# Modulating Nucleosomal H3 Tail Dynamics with Lysine and Serine Modifications

**DOI:** 10.64898/2026.06.30.735535

**Authors:** Brandon J. Adkins, Paul F. W. Sidlowski, Christine E. Jennings, Emma A. Morrison

## Abstract

Nuclear organization is dynamic and originates from the fundamental subunit of chromatin, the nucleosome. Post-translational modification of nucleosomal histones, particularly within intrinsically disordered histone tail regions, provides a dynamic regulatory mechanism of accessibility for chromatin-templated processes. While the epigenomic impacts of lysine acetylation and serine phosphorylation in the histone H3 tail are well-known, how these charge-altering post-translational modifications (PTMs) alter nucleosomal tail conformational dynamics remains incompletely characterized. Given that the functional implications of these PTMs are, at least in part, a consequence of modified nucleosome conformation, systematically cataloging the impact of histone PTMs on nucleosome dynamics provides crucial insight into both baseline cellular activity and epigenetic dysregulation that occurs in disease. Previously, our lab demonstrated that arginine citrullination mimetics lead to regional increases in H3 tail dynamics within nucleosome core particles. Here, we performed nuclear magnetic resonance spin relaxation experiments to investigate the effects of lysine acetylation and serine phosphorylation on H3 tail picosecond-nanosecond (ps-ns) dynamics. Using lysine-to-glutamine and serine-to-glutamate mutations as acetyllysine and phosphoserine mimetics, respectively, we found that these PTMs increase ps-ns conformational dynamics regionally around the PTM site, with a position-dependent effect. Additionally, we show that the type of PTM influences the extent of these increases: in general, the effect of mimetics trends in the order of phosphorylation ≤ acetylation < citrullination, suggesting a tunable method for altering histone tail dynamics. Taken together, these results illustrate the role of nucleosome conformational dynamics in conveying the effects of epigenomic PTMs, elucidating a mechanism of the histone language.

**STATEMENT OF SIGNIFICANCE:** Dynamics are important in expressing the histone language of post-translational modifications. Here, we find that the type and position of a charge-modulating histone tail modification determine the breadth and magnitude of increases in nucleosomal tail dynamics, suggesting a tunable means of perturbing histone tail accessibility and highlighting communication between modifications.

## INTRODUCTION

Dynamic epigenomic regulation is key for proper chromatin organization and regulation of chromatin-templated processes. The fundamental subunit of chromatin is the nucleosome core particle (NCP), an octamer of histone proteins – two each of H2A, H2B, H3, and H4 – wrapped with approximately 147 base pairs of double-stranded DNA (Luger et al., 1997). In addition to a helical core domain, the four canonical histones each contain an intrinsically disordered tail (two in the case of H2A) that protrudes from the core. Positively charged lysine and arginine residues within these histone tails drive intra-and inter-nucleosomal interactions with DNA (Pepenella et al., 2013). Current models describe histone tails as forming “fuzzy” complexes (Fuxreiter, 2018; Tompa & Fuxreiter, 2008) with DNA, not rigidly constrained to a single structure but dynamically bound to nucleosomal DNA in a broad ensemble of conformations (Ghoneim et al., 2021; Jennings et al., 2023; Morrison et al., 2018; Peng et al., 2021b; Rabdano et al., 2021; Stützer et al., 2016; Tsunaka et al., 2022). The tail lysines and arginines, along with serines, are common sites of consequential post-translational modifications (PTMs).

The “language” of histone PTMs describes the indirect and direct epigenomic bases for complex nuclear processes, which originate at the level of the nucleosome (Jenuwein & Allis, 2001; Lee et al., 2010; Strahl & Allis, 2000; Taverna et al., 2007; Winter & Fischle, 2010; Zhao et al., 2021). Histone lysine acetylation, which neutralizes the amino group of the side chain, is generally associated with transcriptionally active euchromatin (Millán-Zambrano et al., 2022). Conversely, histone serine phosphorylation introduces a negative charge and facilitates both gene activation during interphase (Kim et al., 2008; Wang et al., 2023; Yamamoto et al., 2003) and chromatin condensation during mitosis and meiosis (Rossetto et al., 2012; Strahl & Allis, 2000; Wei et al., 1999). Readout of these marks by chromatin-associated proteins is the quintessential indirect mechanism of histone PTMs (Taverna et al., 2007). However, charge-altering histone PTMs, including lysine acetylation, serine phosphorylation, and arginine citrullination, can also directly modulate histone tail-DNA interactions and regulate gene expression by modifying the accessibility of underlying chromatin to effector proteins, higher-order chromatin interactions, and the stability and conformation of nucleosomes (Bowman & Poirier, 2015; Millán-Zambrano et al., 2022; Peng et al., 2021a).

While readout of histone PTMs has been extensively studied (Taverna et al., 2007), the direct effects conveyed through protein conformational dynamics remain incompletely understood. Unmodified nucleosomal histone tails assume a broad ensemble of DNA-bound states, with dynamic motions on the picosecond (ps) to millisecond timescales (Furukawa et al., 2020; Ghoneim et al., 2021; Jennings et al., 2023; Lehmann et al., 2020; Morrison et al., 2021, 2018; Stützer et al., 2016; Tsunaka et al., 2022; Zandian et al., 2021). These baseline tail dynamics are altered in numerous nucleosomal contexts, including the incorporation of H1 linker histone (Stützer et al., 2016), the presence of linker DNA (Furukawa et al., 2020), and changes in nucleosome assembly state (Morrison et al., 2021). Furthermore, some charge-modifying PTMs, such as H3 Lys14-aceylation or Ser(10/)28-phosphorylation, modulate tail-DNA interactions, leading to increases in H3 tail dynamics (Pelaz et al., 2020; Stützer et al., 2016), and phosphoserine has the potential to introduce tail-tail interactions (Bailey et al., 2013; Papamokos et al., 2012; Zheng & Hayes, 2003). These changes in dynamics are implicated in NCP accessibility to histone reader, writer, and eraser proteins (Gatchalian et al., 2017; Morrison et al., 2018; Stützer et al., 2016) and influence the rate of further enzymatic modification (Furukawa et al., 2020; Kowalczyk & Morrison, 2026; Stützer et al., 2016), providing a crucial role for histone dynamics in normal cellular functioning.

We seek to further elucidate the role of nucleosome dynamics in the histone language by systematically and comprehensively characterizing the impact of site-specific PTMs on tail dynamics via nuclear magnetic resonance (NMR) spectroscopy. At approximately 200 kDa, NCPs exceed the conventional molecular-weight limit for backbone-amide NMR detection (Foster et al., 2007; Kleckner & Foster, 2011; Ruschak & Kay, 2009; Schütz & Sprangers, 2020). However, even in fully formed NCPs, the high mobility of the intrinsically disordered histone tails generates observable cross peaks. Accordingly, histone tail dynamics have been studied by multiple groups, including ours, via NMR nuclear spin relaxation (NSR) experiments (Fuchs et al., 2026; Furukawa et al., 2022; Hammonds et al., 2026; Jennings et al., 2023; Kim et al., 2023; Morrison et al., 2021, 2018; Pelaz et al., 2020; Rabdano et al., 2021; Stützer et al., 2016; Zandian et al., 2021; Zhou et al., 2012). NSR observables (R_1_, R_2_, and hnNOE) are influenced by minor magnetic field fluctuations that result from fast-timescale, internal protein motions, and thus, report on ps-to-nanosecond (ns) conformational dynamics with residue resolution. Monitoring changes in NSR observables in response to alterations to a histone (e.g., PTMs) informs our understanding of nucleosome conformational dynamics and their perturbations.

As the longest of the histone tails, the H3 N-terminus contains four arginines, six lysines, and two serines that are biologically relevant sites of PTM, positioning these residues as appealing candidates for conformational dynamics studies. Basic residues within the tail form dynamic points of interaction with DNA that are segregated by two flexible TGG motifs, the second of which is part of a larger “hinge” region (S28-K36) (Furukawa et al., 2020; Morrison et al., 2021; Stützer et al., 2016). Previously, we investigated the effect of arginine neutralization (as a mimetic for citrullination) on H3 tail dynamics (Jennings et al., 2023). Neutralization of all four H3 tail arginines via glutamine mutation disrupted tail-DNA interactions and led to a global increase in dynamics, whereas the effects of individual arginine neutralizations were regional and site-dependent. Here, we use NMR NSR experiments to expand upon this relationship between histone tail PTMs and nucleosome dynamics, investigating the effects of lysine acetylation and serine phosphorylation on H3 tail dynamics at residue-level resolution. In our prior arginine citrullination study, we reported baseline ps-ns dynamics across the H3 tail (residues T3-K36) in unmodified NCPs (Jennings et al., 2023). These wild-type (WT) values will again serve as a comparative basis for the H3 lysine-to-glutamine (KQ) and serine-to-glutamate (SE) NCPs examined in this study. Similar to arginine neutralization, individual mimetics of lysine acetylation and serine phosphorylation lead to regional increases in H3 tail dynamics. For residues in proximity to each other (e.g., R26Q, K27Q, and S28E), this increase was greatest for individual arginine mutations, followed by lysine, and then serine. The cumulative effect from the neutralization of all six lysines, however, is larger than that from all four arginines, highlighting the importance of both type and number of PTM(s) on histone dynamics. Changes in H3 tail dynamics have a positional dependence, and mutations adjacent to the “hinge” generally had the greatest impact on dynamics. These results provide insight into the mechanistic underpinnings of histone PTMs and PTM crosstalk by detailing fundamental changes in histone tail motions that are implicated in chromatin-templated processes.

## RESULTS

### Site-specific incorporation of lysine acetylation mimetics alters the H3 tail conformational ensemble

To catalog the effects of lysine acetylation on the H3 tail conformational ensemble and dynamics, NCPs were reconstituted with ^15^N-H3 KQ mutants and tested by NMR spectroscopy. The H3 tail contains six lysine residues N-terminal to the hinge that are visible in ^1^H-^15^N backbone spectra of reconstituted NCPs – K4, K9, K14, K18, K23, and K27. While selective acetylation of these individual sites is challenging to achieve enzymatically by histone acetyltransferases, site-specific modification of a single lysine can be mimicked by mutagenesis to glutamine (Wang & Hayes, 2008), which yields the critical side-chain neutralization. ^1^H-^15^N HSQC and NSR experiments were conducted for six single-KQ (e.g., K4Q-H3, etc.) NCPs and one sextuple-KQ (i.e., K4/9/14/18/23/27Q-H3; henceforth, 6xKQ-H3) NCP (**Figures S1-S4)**. In the resulting spectra, most H3 tail residues, including the sites of KQ mutation, display a single resonance, but doublet peaks are observed for some NCPs (**Figure S1; Table S1)**. Doublet peaks have been observed in past NMR studies of the H3 tail in the context of NCPs (Jennings et al., 2023; Morrison et al., 2021) and may result from differences in the conformational ensembles of the two H3 tails due to asymmetry in the Widom 601 DNA sequence or an effect from slower timescale dynamics. In general, the residues displaying doublet peaks in this study align with those we previously reported for arginine-to-glutamine (RQ) H3-NCPs (**Table S1**) (Jennings et al., 2023).

CSPs between ^1^H-^15^N HSQC spectra of WT- and KQ-H3-NCPs are sensitive probes of residue-level differences in the H3 tail conformational ensemble. For all single-KQ-H3-NCPs examined in this study, the largest CSPs are localized to the mutated residue and its neighbors (**Figure 1**). As with single-RQ-H3-NCPs (Jennings et al., 2023), non-mutated residues show modest perturbations (CSPs < 0.06 ppm), with the notable exception of T22 (0.15 ppm) in the K23Q-H3-NCP. While CSPs greater than 0.04 ppm do not occur more than five residues away from the point of mutation, small CSPs are observed remote from the mutation, as exemplified by residues A24-A29 in ^15^N-K4Q-H3-NCP (**Figure 1**). Similar low-magnitude distal CSPs have been observed with other charge-modulating H3 tail mutations (Fuchs et al., 2026; Jennings et al., 2023; Pelaz et al., 2020) and suggest that lysine neutralizations alter the structural ensemble of the H3 tail even at positions that cross the first TGG motif.

**Figure 1.**
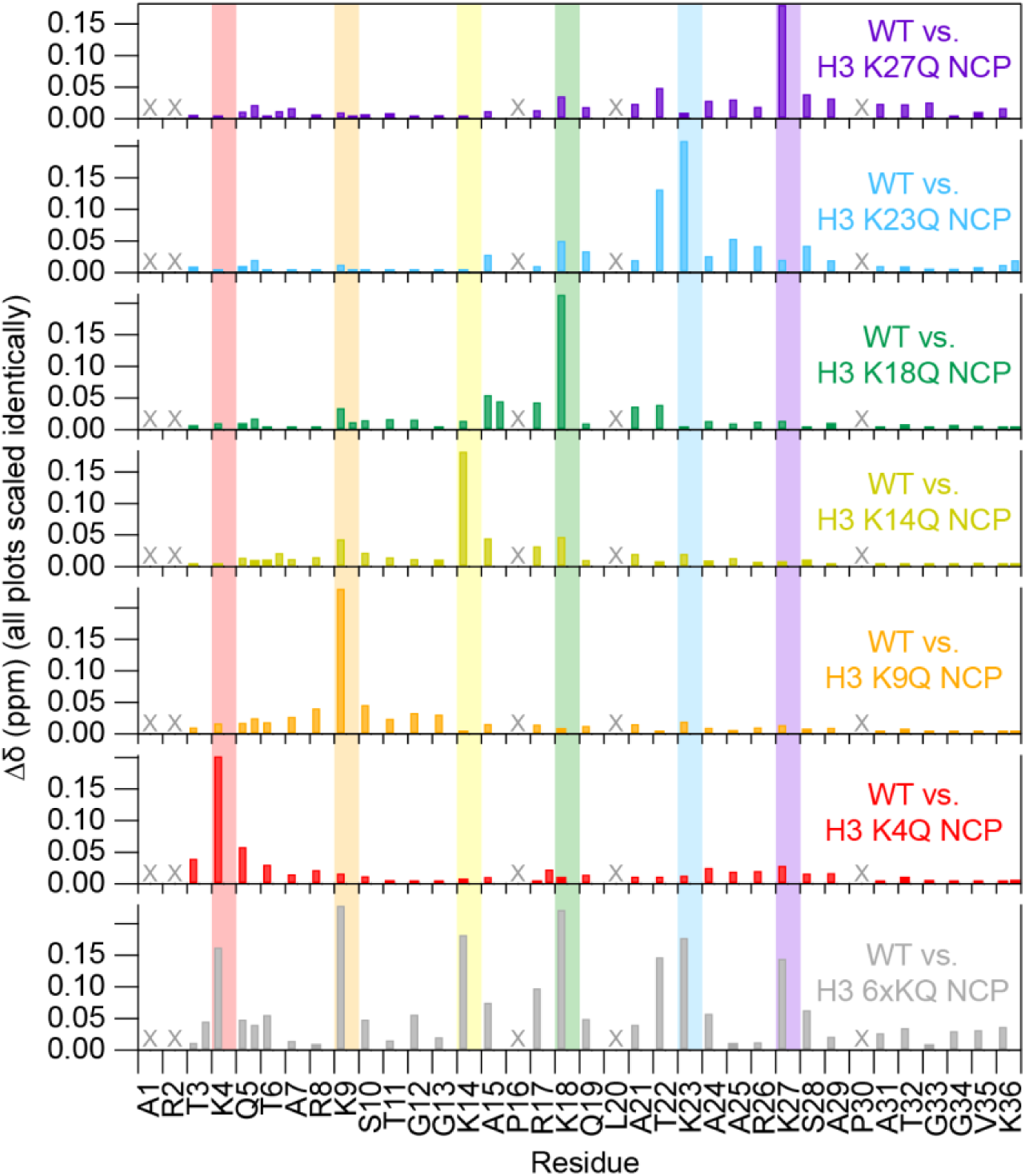
Chemical shift perturbations by lysine neutralizations reveal largely small, localized conformational changes in the H3 tail. Chemical shift perturbations (Δδ) between WT and each mutant are plotted as a function of H3 tail residue for ^1^H-^15^N HSQC spectra collected at 800 MHz and 31°C, with 0 mM added salt. Two values are plotted for doublet peaks, and “X” symbols indicate residues omitted from analysis. The y-axes for each mutant are scaled identically and consistent with Figure 3.

### Neutralizing all six H3 tail lysines results in a global increase in nucleosomal H3 tail mobility

To assess differences in the ps-ns conformational dynamics of KQ- versus WT-H3-NCPs, we performed NMR NSR experiments. When comparing mutants to WT-H3-NCP, an observed decrease in R_2_/R_1_ ratio and hnNOE value corresponds to an increase in ps-ns timescale dynamics at a given position (Kleckner & Foster, 2011). We previously determined that the residue-averaged longitudinal spin relaxation rate (R_1_) across the H3 tail of WT-H3-NCP is 1.07 ± 0.09 s^−1^ while the average transverse spin relaxation rate (R_2_) is 22 ± 8 s^−1^, resulting in an average R_2_/R_1_ ratio of 21 ± 9 (**Figure 2A; Table S2**) (Jennings et al., 2023). The residue-averaged H3 tail hnNOE value in WT-H3-NCP is 0.35 ± 0.12 (**Figure 2B; Table S2**) (Jennings et al., 2023).

**Figure 2.**
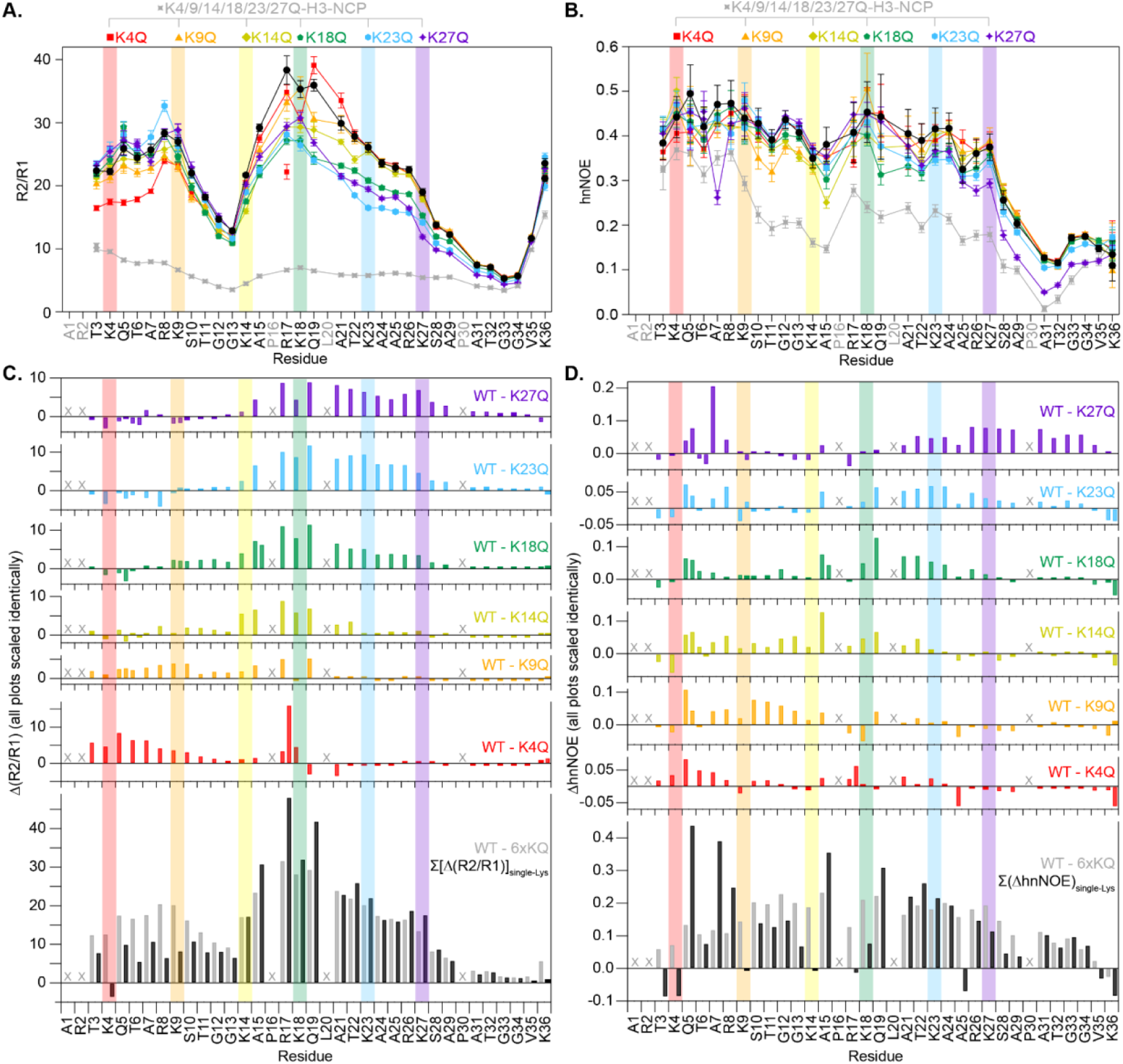
Acetyllysine mimetics increase nucleosomal H3 tail dynamics regionally. R_2_/R_1_ ratio (**A**), hnNOE (**B**), Δ(R_2_/R_1_) (**C**), and ΔhnNOE (**D**) are plotted as a function of H3 tail residue for unmodified NCP (black), 6xKQ-H3-NCP (gray), and single-KQ-mutant-H3-NCPs (rainbow colors, as labeled). For R_2_/R_1_ (**A**) and hnNOE (**B**), the error bars are determined from standard error propagation of either the error from the covariance matrix in fitting R_1_ and R_2_ or the spectral noise in the hnNOE spectra. The difference plots (**C**, **D**) are calculated between WT and each mutant. The lower plots in (**C**) and (**D**) show the sum of the Δ(R_2_/R_1_) or ΔhnNOE (excluding doublets) for six individual KQ-H3-NCPs (black). “X” symbols indicate residues that were not included in analysis. Two values are plotted for doublet peaks (see Materials and Methods). Y-axes within (**C**) and (**D**) are scaled identically and consistent with Figure 4. Data were collected in the absence of added salt at 800 MHz and 31°C.

Of all KQ-H3-NCPs examined in this study, the 6xKQ-H3-NCP displays both the largest and broadest decreases in per-residue profiles of R_2_/R_1_ ratios and hnNOE values, indicative of global increases in H3 tail dynamics (**Figure 2**). The neutralization of all six H3 tail lysines results in R_2_/R_1_ ratios ranging from 3.4 (G33) to 15.4 (K36), with an average value of 6 ± 2 – a dramatic decrease of 66% from WT-H3-NCP (**Table S2**). The hnNOE values decrease an average of 40% to 0.21 ± 0.09, ranging from 0.01 (A31) to 0.37 (K4Q) (**Table S2**). While both observables retain local minima near the two flexible TGG motifs, the H3 tail is more uniformly dynamic in this “hyperacetylated” state as compared to WT-H3-NCP.

Of note, there are regional discrepancies between trends in the R_2_/R_1_ ratios and hnNOE values, which hold true for nearly all NCPs tested in this study. A probable explanation is twofold. First, hnNOE experiments are of lower sensitivity and, therefore, result in noisier spectra. Second, the R_2_/R_1_ ratio is more strongly influenced by contributions from overall protein tumbling than hnNOE values. Consistent among both data plots, however, is a global increase in ps-ns dynamics across the 6xKQ-H3 tail, presumably due to the disruption of electrostatic interactions between tail lysines and nucleosomal DNA. While the four unmodified tail arginines are still present to contact DNA, they are clearly not sufficient to completely preserve regional distinctions in H3 tail ps-ns dynamics with all six lysines neutralized.

### Neutralizing individual H3 tail lysines increases nucleosomal H3 tail mobility regionally

To determine the positional effects of lysine neutralization, the same NSR experiments were performed on six NCPs containing individual KQ mutations within the H3 tail (**Figure 2A, 2B; Table S2**). Differences in the R_2_/R_1_ and hnNOE profiles for each single KQ mutant as compared to WT [i.e., Δ(R_2_/R_1_) and ΔhnNOE, respectively] were used to assess the impact of site-directed lysine neutralization on ps-ns tail dynamics (**Figure 2C, 2D; Table S3**). As with the 6xKQ-H3-NCP, neutralization of single tail lysines primarily increases H3 tail dynamics, but in a regional rather than global manner. Profiles of Δ(R_2_/R_1_) and ΔhnNOE as a function of residue reveal that the largest changes in dynamics generally peak near the site of KQ mutation in single KQ-H3-NCPs but are not simply localized near the mutation. (Similar to observations with R26Q-H3-NCP(Jennings et al., 2023), perturbations of R_2_/R_1_ are likely skewed to the N-terminal side of the mutation for K27Q- and K23Q-H3-NCP due to overall particle tumbling dominating at positions closer to the nucleosomal core.) The regional change in dynamics is further illustrated by the breadth of consecutive H3 tail residues having non-overlapping error bars between the KQ- and WT-H3 R_2_/R_1_ ratios and hnNOE values (**Figure 2C, 2D; Table 1**). Importantly, the effect of single KQ neutralizations extends to include 1-3 other lysine positions.

**Table 1.** Compilation of the breadth of the effect of charge-modulating lysine, serine, and arginine mutations on H3 tail dynamics within mutant-H3-NCPs as compared to WT-H3-NCP. The H3 tail region affected by lysine, serine, or arginine mutation was defined by a string of at least three residues with non-overlapping error bars (with possible interruption by a single residue with overlapping error bars). The range was determined for each mutant from either R_2_/R_1_ or hnNOE data sets. The boundaries of the region are listed along with the number of residues within this range for facile assessment of the effect. Data from arginine-to-glutamine mutations are from Jennings et al., 2023.

| Mutant | $R_2/R_1$ | hnNOE |
| --- | --- | --- |
| K4Q | T3-A21 (19 residue range) | Q5 and A25 (1 residue range) |
| K9Q | T3-Q19 (17 residue range) | S10-G13 (4 residue range) |
| K14Q | R8-T22 (15 residue range) | G12-A15 (4 residue range) |
| K18Q | K9-A29 (21 residue range) | Q19-T22 (4 residue range) |
| K23Q | G12-K36 (25 residue range) | T22-R26 (5 residue range) |
| K27Q | K14-K36 (23 residue range) | T22-V35 (14 residue range) |
| 6xKQ | T3-K36 (34 residue range) | K4-V35 (32 residue range) |
| Mutant | $R_2/R_1$ | hnNOE |
| S10E | T3-T22 (20 residue range) | T11-A15 (5 residue range) |
| S28E | K14-K36 (23 residue range) | T22-K36 (15 residue range) |
| 2xSE | T3-K36 (34 residue range) | R26-G34 (9 residue range) |
| Mutant | $R_2/R_1$ | hnNOE |
| R2Q | T3-G12 (10 residue range) | T3-T5 (3 residue range) |
| R8Q | T3-T22 (20 residue range) | A7-A15 (9 residue range) |
| R17Q | R8-A29 (22 residue range) | K14-T22 (9 residue range) |
| R26Q | G13-V35 (23 residue range) | T22-V35 (14 residue range) |
| R2/8/17/26Q | T3-K36 (34 residue range) | T3-G34 (32 residue range) |

There is a positional dependence to the breadth and depth of the effect of single lysine neutralizations. The K27Q- and K23Q-H3 mutations have the broadest impact on hnNOE values and R_2_/R_1_ ratios, respectively (**Table 1**). In general, proximity of a lysine neutralization to the nucleosomal core correlates with the sum of ΔhnNOE values (Σ_ΔhnNOE_) (**Table S3**). The sums rank as K27Q (1.0) > K18Q (0.7) > K14Q (0.6) ≈ K23Q (0.6) > K9Q (0.4) > K4Q (0.2) or K27Q (1.3) > K23Q (1.0) ≥ K14Q (0.9) ≈ K9Q (0.9) ≈ K18Q (0.9) > K4Q (0.6), depending on whether raw or absolute values are used, respectively. [Summation of absolute values better captures changes in either direction (see below) but also accentuates noise.] Additionally, the sum of Δ(R_2_/R_1_) ratios (Σ_Δ(R2/R1)_) confirms that increases in ps-ns dynamics are greatest for the KQ mutations that lie between the two TGG motifs. Depending on whether raw or absolute values are used, ranking by Σ_Δ(R2/R1)_ is K23Q (94) > K18Q (90) > K27Q (77) >> K14Q (57) > K4Q (53) > K9Q (44) or K23Q (120) > K27Q (102) ≥ K18Q (97) >> K4Q (71) > K14Q (60) > K9Q (48), respectively. Considering that effects on R_2_/R_1_ are expected to be muted towards the core, we speculate that the K27Q neutralization had the largest impact on H3 tail ps-ns dynamics within the lysine neutralization series, similar to R26Q within the arginine neutralization series (Jennings et al., 2023). This position is adjacent to the flexible H3 tail hinge region, and neutralization extends the mobile hinge further toward the N-terminus.

Interestingly, comparison of R_2_/R_1_ ratios suggests the possibility that neutralizations C-terminal to the first TGG motif (i.e., K18Q, K23Q, and K27Q) decrease the dynamics of multiple residues preceding this TGG motif. This trend was also observed in data collected on R26Q-H3-NCP (Jennings et al., 2023). This results in a discrepancy between the Δ(R_2_/R_1_) for the 6xKQ-H3-NCP versus the residue-by-residue sum of the Δ(R_2_/R_1_) for the six individual KQ mutants from residue T3 to G13 (**Figure 2C**, gray versus black bars). Similarly, mutations that occur N-terminal to the first TGG motif may decrease the dynamics of many succeeding residues, though to a lesser extent. While present, this phenomenon is not as strongly recapitulated in the ΔhnNOE data set. These observations are consistent with a model in which the tail is divided into regions separated by the TGG motifs, and neutralization in one region may create stronger contacts between DNA and the basic residues in the other region. The induced relative rigidity is overcome when all six lysines are neutralized.

### Site-specific incorporation of phosphoserine mimetics affects the conformational ensemble of nucleosomal H3 tails

In addition to lysine acetylation, serine phosphorylation is another critical modification in the histone language. Rather than neutralizing a cationic side chain, phosphorylation introduces a negative charge, weakening tail-DNA interactions and introducing the potential for intra- or inter-tail interactions with basic residues. There are two serines in the H3 tail, each of which is routinely modified in various biological contexts (Rossetto et al., 2012; Strahl & Allis, 2000; Wei et al., 1999). To determine the impact of phosphorylation of these sites on H3 tail conformation and dynamics, we reconstituted NCPs with ^15^N-H3 containing either individual (S10E- or S28E-H3) or dual (S10/28E-H3; henceforth, 2xSE-H3) serine-to-glutamate (SE) phosphomimetics. Like KQ-H3-NCPs, these SE-H3-NCPs were each subjected to four sets of NMR experiments: ^1^H-^15^N HSQCs, ^15^N R_1_ relaxation series, ^15^N R_2_ relaxation series, and hnNOE (**Figures S5-S8)**. Similar to KQ- and RQ-H3-NCPs, H3 tail CSPs resulting from individual serine phosphomimetics were small (CSPs < 0.06 ppm) for non-mutated residues, with the exception of T11 for S10E-H3-NCP and A29 and T32 for S28E-H3-NCP (**Figure 3**). The largest CSPs again localized to residues immediately surrounding the site of mutation, with all residues more than four positions away having CSPs less than 0.04 ppm. Small distal CSPs are noted, consistent with the KQ data and a previous study with S28Q-H3-NCP (Pelaz et al., 2020).

**Figure 3.**
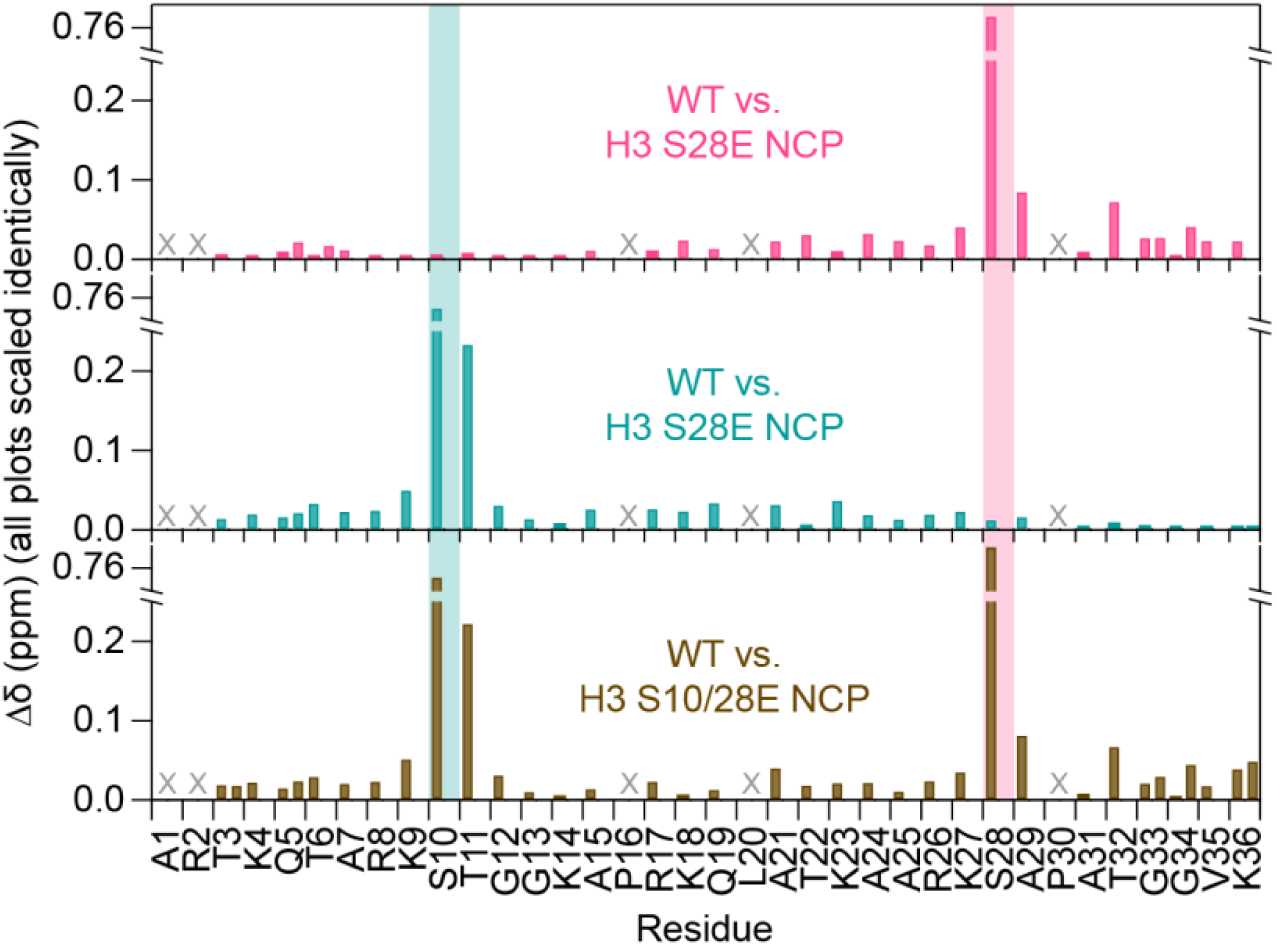
Chemical shift perturbations by phosphoserine mimetics indicate predominantly localized conformational changes in the H3 tail. Chemical shift perturbations (Δδ) between WT and each mutant are plotted as a function of H3 tail residue. ^1^H-^15^N HSQC spectra were collected at 800 MHz and 31°C, with 0 mM added salt. Two values are plotted for doublet peaks. “X” symbols indicate residues omitted from analysis. The y-axes for each mutant are scaled identically and consistent with Figure 1.

### Phosphomimetics of both H3 tail serines increase nucleosomal H3 tail mobility but retain restrictions in motion

NSR experiments were used to quantify the effect of SE phosphomimetics on H3 tail ps-ns dynamics. Complete serine “phosphorylation” in the 2xSE-H3-NCP primarily increases H3 tail dynamics, as indicated by a 20% decrease in the residue-averaged R_2_/R_1_ ratio (17 ± 6) relative to WT-H3-NCP, with individual residues ranging from 5 (G33) to 27 (K18) (**Figure 4A; Table S2**). This is corroborated by a 7.5% decrease in the average hnNOE value (0.3 ± 0.1), which varies from 0.07 (A31) to 0.46 (K4) (**Figure 4B; Table S2**). This increase in H3 tail dynamics is not as extensive as that observed in the 6xKQ-H3-NCP, at least in part because there are fewer sites of modification. Normalizing the percent change in R_2_/R_1_ ratio and hnNOE value for these two NCPs on a per-mutation basis suggests that the impact per acetyllysine mimetic (11% and 7%, respectively) is similar to that of an individual phosphomimetic (10% and 4%, respectively).

**Figure 4.**
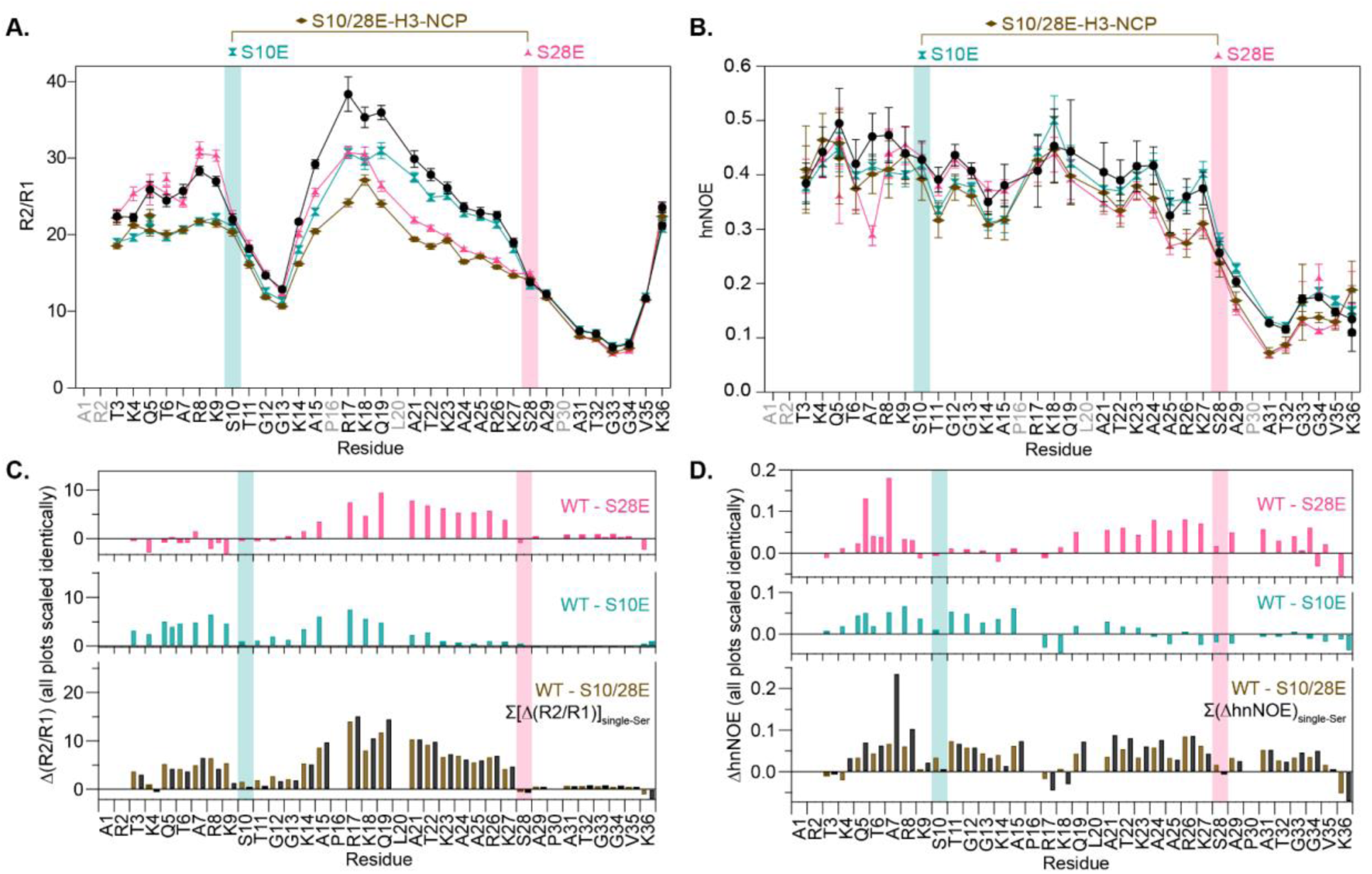
Serine phosphomimetics cause a regional increase in nucleosomal H3 tail dynamics. R_2_/R_1_ ratio (**A**), hnNOE (**B**), Δ(R_2_/R_1_) (**C**), and ΔhnNOE (**D**) are plotted as a function of H3 tail residue for WT-, 2xSE-, and single-SE-mutant-H3-NCPs (colors as labeled). For R_2_/R_1_ (**A**) and hnNOE (**B**), the error bars are determined from standard error propagation of either the error from the covariance matrix in fitting R_1_ and R_2_ or the spectral noise in the hnNOE spectra. The difference plots (**C**, **D**) are calculated between WT and each mutant. The lower plots in (**C**) and (**D**) show the sum of the Δ(R_2_/R_1_) or ΔhnNOE for two individual SE-H3-NCPs (black). “X” symbols indicate residues that were omitted from analysis. Two values are plotted for doublet peaks (see Materials and Methods). Y-axes within (**C**) and (**D**) are scaled identically and consistent with Figure 2. Data were collected at 800 MHz and 31°C, in the absence of added salt.

Unlike complete lysine neutralization in the H3 tail, the 2xSE-H3-NCP preserves regional distinctions in dynamics, which results in larger variations in R_2_/R_1_ ratios and hnNOE values across tail residues for the 2xSE- versus 6xKQ-H3 samples. In both NCPs, the hinge region persists as the most dynamic portion of the H3 tail. The elevated flexibility of the H3 hinge region relative to the rest of the tail is less pronounced for the 6xKQ-H3- NCP, especially in the R_2_/R_1_ curve (**Figure 2A**) but remains readily apparent in the 2xSE-H3-NCP (**Figure 4A**).

### Individual serine phosphomimetics increase regional nucleosomal H3 tail mobility

Lastly, we tested the effects of single-site serine phosphomimetics on H3 tail dynamics (**Figure 4A, 4B; Table S2**). Similar to single basic-residue neutralizations, Δ(R_2_/R_1_) and ΔhnNOE profiles highlight a regional effect for phosphomimetics on nucleosomal H3 tail dynamics (**Figure 4C, 4D; Table S3**). In considering the magnitude and breadth of effect, the Σ_Δ(R2/R1)_ (when absolute values are used), Σ_ΔhnNOE_, and range of perturbed residues for SE-H3-NCPs (**Figure 5**; **Tables 1, S3**) are consistent with observations on KQ- and RQ-H3-NCPs and suggest that the closer a given PTM occurs to the H3 hinge region, the greater its impact on dynamics. Notably, we were unable to reproduce the global effect of the S28E mutation on tail dynamics observed by Pelaz et al., who reported that the mobility of the entire H3 tail increased by ∼20% (Pelaz et al., 2020).

**Figure 5.**
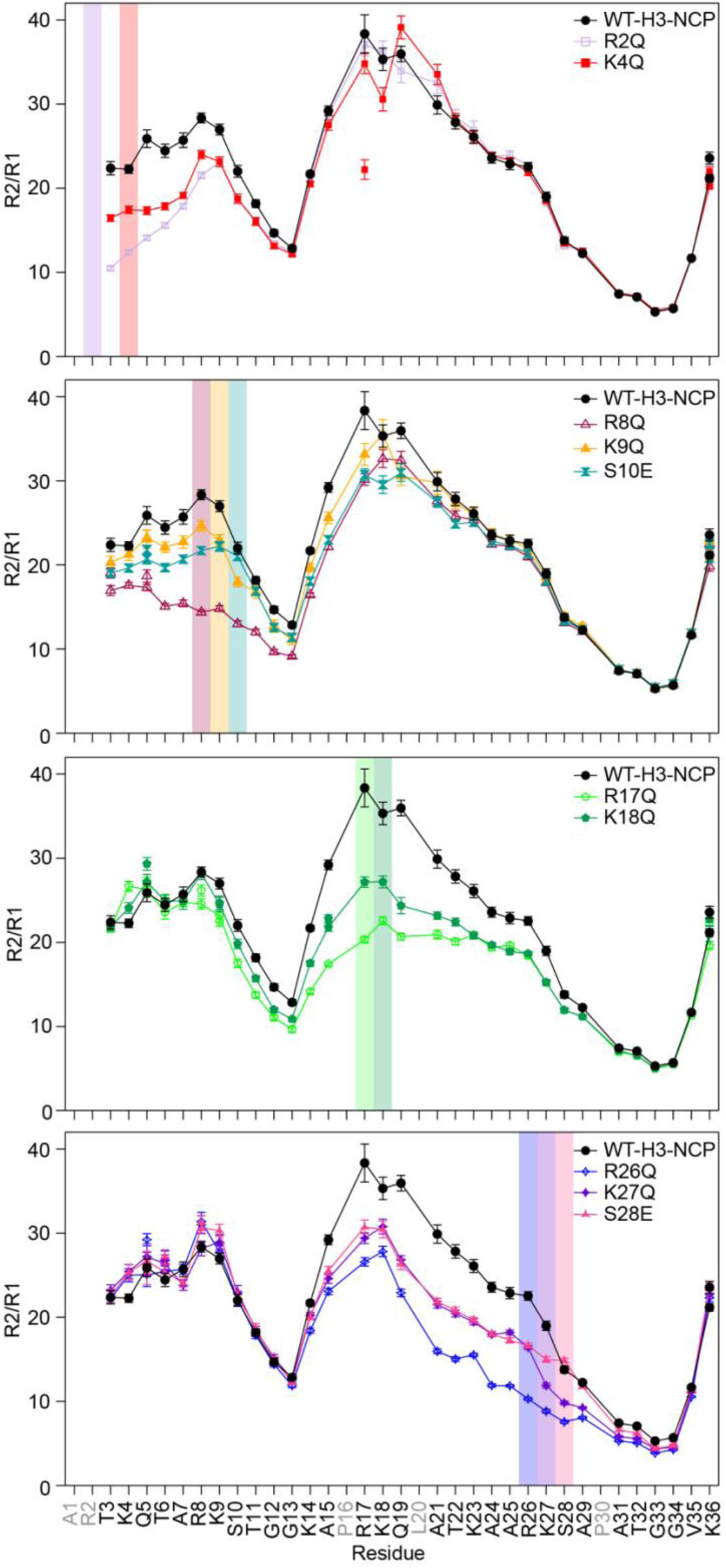
Direct comparison of H3 tail charge-modulating PTM mimetics highlights differences in magnitude and breadth of effect. Mutated positions are marked by a colored background bar. Error bars represent standard error propagation from the covariance matrix in fitting R_1_ and R_2_ decay curves. Note that some error bars are smaller than the symbols. Residues without data plotted for any sample are colored gray in the x-axis labels. Data are from Figures 2 and **4** and Jennings et al., 2023 and were collected in the absence of added salt at 800 MHz and 31°C.

Surprisingly, the singly “phosphorylated” residues themselves display R_2_/R_1_ ratios and hnNOE values that are unusually consistent with WT-H3 dynamics (**Figure 4**). This is in contrast to individual KQ- and RQ-H3-NCPs, for which the site of mutation is among the residues with the largest Δ(R_2_/R_1_) and ΔhnNOE values (**Figures 2, 5, S9**). For example, when comparing the impact of the S28 phosphomimetic to the neighboring acetyllysine mimetic, we observe that S28E-H3-NCP yields similar R_2_/R_1_ and hnNOE curves to K27Q-H3-NCP, deviating only near the site of mutation (K27-A29) (**Figures 5**, **S9**).

Crosstalk across the first TGG motif is also observed with the phosphomimetics. Out of all NCPs tested in this study, S10E-H3 is the individual mutation that most effectively transmits increases in tail dynamics across the first TGG motif, as evidenced by the Δ(R_2_/R_1_) between T3 (the first NMR visible residue) and T22, and is similar to R8Q-H3 in this regard (**Figures 5, S9; Table 1**) (Jennings et al., 2023). The N-terminal portion of the S28E-H3 tail (preceding the first TGG motif) contains six residues with negative R_2_/R_1_ values, which is consistent with observations made upon neutralizing residues between the two TGG motifs (i.e., K18Q, K23Q, R26, and K27Q) and further supports that increased dynamics in the central region of the tail may lead to decreased dynamics in the N-terminal region.

## DISCUSSION

In this study, we used NMR spectroscopy to quantify the effects of acetyllysine and phosphoserine mimetics on nucleosomal histone H3 tail conformation and fast-timescale (ps-ns) dynamics. We demonstrate that the disruption of tail anchor points through the neutralization of basic residues (via lysine-to-glutamine mutations) or the introduction of repulsive forces (via serine-to-glutamate mutations) increases H3 tail ps-ns dynamics. We also catalog the regional, position-dependent effects of individual residue perturbations. Overall, our results support that, like arginine citrullination (Jennings et al., 2023), lysine acetylation and serine phosphorylation primarily increase the conformational dynamics of the H3 tail, with implications for accessibility to readers, writers, and erasers of the histone tails and crosstalk between PTMs.

For a given PTM mimetic, mutation of the maximal number of sites (i.e., 6xKQ-H3 and 2xSE-H3) yields global increases in tail mobility. Such NCPs represent H3 tail lysine hyperacetylation and total serine phosphorylation, both of which are associated with active gene expression (Clayton et al., 2006; Davie & Chadee, 1998; Lempiäinen & Garcia, 2023; Rossetto et al., 2012; Sun et al., 2007). The increases in dynamics associated with these epigenetic states serve as a fundamental mechanism for this regulation by loosening chromatin structure and providing crucial accessibility to transcriptional machinery and/or chromatin remodelers (Bowman & Poirier, 2015; Chatterjee et al., 2011; Millán-Zambrano et al., 2022). Our results are consistent with NMR dynamics studies of nucleosomes enzymatically acetylated by Gcn5 or phosphorylated by Aurora B (Stützer et al., 2016).

We found that single-site acetyllysine and phosphoserine mimetics influence H3 tail dynamics in a regional (rather than global) and site-dependent manner. Consistent with the model from our arginine neutralization study (Jennings et al., 2023), this is most pronounced for mutations near the H3 tail hinge region, such as K27Q-H3 and S28E-H3. These mutations increase ps-ns dynamics across ∼39%-64% (depending on determination by hnNOE or R2/R1) of the tail by weakening the R26/K27 anchor points and extending the flexible hinge N-terminally. This model also aligns with the epigenomic impacts of H3 K27ac and H3 S28ph, which are both marks of transcriptional activation (Konuma & Zhou, 2024; Lau & Cheung, 2011; Sun et al., 2007). Conversely, the more N-terminal mutations generally made the smallest impacts while still influencing more than a third of the tail (as determined by R2/R1). In addition, although the magnitudes are small, the breadth of impact (as determined by R2/R1) crosses the first TGG motif for charge perturbations N- and C-terminal to the motif. Interestingly, the data support the possibility that some mutations have the potential to constrain the dynamics of remote residues across the first TGG motif from the mutation. Overall, the breadth of impact of the charge-altering mutations in this study is expected to be at least as broad in the presence of physiologically relevant salt concentrations, as observed for arginine neutralizations (Jennings et al., 2023).

When comparing H3 tail PTM mimetics, a dynamics hierarchy is apparent. For single-site mutations of neighboring lysines, serines, and arginines, the arginine generally has the largest impact in terms of breadth and magnitude of effect, followed by lysine and serine. However, the cumulative effect of neutralizing all six tail lysines exceeds that of neutralizing the four tail arginines, highlighting the importance of both the type and number of PTMs. Our findings are in agreement with simulation data, which show that H3 tail arginines form both a greater number of and more energetically favorable interactions with DNA than do tail lysines (Li & Kono, 2016; Morrison et al., 2018; Peng et al., 2021b). Compared to adjacent RQ and KQ mutations, the glutamate backbone amides in S10E-and S28E-H3-NCPs are more restricted, displaying R_2_/R_1_ ratios and hnNOE values that closely align those of the respective serines in WT-H3-NCPs. Nevertheless, neighboring residues are more dynamic than in the WT-H3-NCP, but with a muted increase in dynamics extending 1-2 residues on each side of the mutation (from the hnNOE and R2/R1 profiles; **Figures 5, S9**). We speculate that this local restriction in motion results from the negatively charged glutamate sidechain interacting with histone tail basic residue(s). Simulations of H3 tail peptides support strong intra-tail interactions for S10E-R8 and S28E-R26 (Papamokos et al., 2012; Preez & Patterton, 2017), but under the conditions of our experiments, there is limited support for a restriction in R8 and R26 dynamics. While there could conceivably be an S10E-K9 or S28E-K27 interaction, the restricted dynamic increase for K9 and K27 is likely the result of simply being adjacent to the restricted S10E and S28E. We favor an inter-tail interaction, perhaps with an H4 tail arginine (Mascotti & Lohman, 1997; Woods & Ferré, 2005). Together, these individual PTM mimetics support a mechanism for fine-tuning the regional dynamics and accessibility of the H3 tail, which may extend to other tails (Furukawa et al., 2020; Kowalczyk & Morrison, 2026).

Our study has several experimental limitations: i) The data here report the impacts of charge-altering histone PTM mimetics on ps-ns dynamics in the H3 tail, which compound to generate longer-timescale motions (Lehmann et al., 2020). While the effective CPMG strength (500 Hz) used in our R2 experiments may not be sufficient to completely suppress contributions from µs-ms dynamics, which could be differentially affected by mutations, we interpret our data as reporting on ps-ns timescale dynamics. Further experiments are necessary to distinguish dynamics across timescales. ii) Our experiments may not be able to distinguish changes in dynamics that reflect residue-level DNA-association/dissociation from changes in the conformation and mobility of the DNA that the tails interact with. iii) Our approach does not distinguish intra- and inter-nucleosomal tail-DNA interactions. We speculate that intra-nucleosomal interactions dominate in the data presented here (collected in the absence of added salt for optimal spectral quality). Similar trends have been observed with 150 mM KCl and within phase-separated condensates, where inter-nucleosomal interactions are presumed to also be present (Hammonds et al., 2026; Jennings et al., 2023). iv) While the synthetic Widom 601 sequence is standard for *in vitro* NCP and nucleosome studies, the dynamics may not be representative of the native genome. However, because Widom 601 DNA stabilizes the nucleosome (Lowary & Widom, 1998), we speculate that the increases in tail dynamics observed in this study would only be amplified if the NCPs were instead reconstituted with a native sequence (Chen et al., 2025). Future studies are necessary to explore the limitations of this study.

Altogether, this work expands and refines an emerging model of dynamics in the histone language. Histone tails and their epigenetic marks are dynamic regulators of nucleosome conformation and stability, interactions with chromatin components (i.e., histones and DNA) and chromatin-associated proteins, and accessibility to these chromatin-associated proteins (Bowman & Poirier, 2015; Millán-Zambrano et al., 2022; Peng et al., 2021a). By systematically characterizing mimetics of H3 tail lysine acetylation and serine phosphorylation, we build upon this body of work. In combination with our study on arginine citrullination mimetics (Jennings et al., 2023), our results highlight that the number, type, and position of the charge-modulating PTM finely tune conformational dynamics. Our studies have important implications for crosstalk between PTMs—the neutralization of a single basic residue or addition of a single negative charge regionally increases the ps-ns dynamics of the tail, with the effect extending over several other key epigenetic regulatory positions (**Figure 6**). The biological implication of this crosstalk is increased accessibility of these other positions for modification, readout, or erasure. Other studies have reported that charge-altering modifications to arginines, lysines, and serines increase binding of readers or writers to other positions on the same tail and subsequent enzymatic modification of the same or other tail (Furukawa et al., 2020; Gatchalian et al., 2017; Morrison et al., 2018; Peng et al., 2021b; Stützer et al., 2016). Functionally relevant PTMs on other canonical and non-canonical histones can alter nucleosome dynamics (Bowman & Poirier, 2015; Furukawa et al., 2020; Kim et al., 2023; Nosella et al., 2024; Stützer et al., 2016) and should be further explored. By comprehensively characterizing the role of conformational dynamics in the histone language, these studies will provide insight into proper genomic regulation that can be leveraged to correct dysfunction in disease.

**Figure 6.**
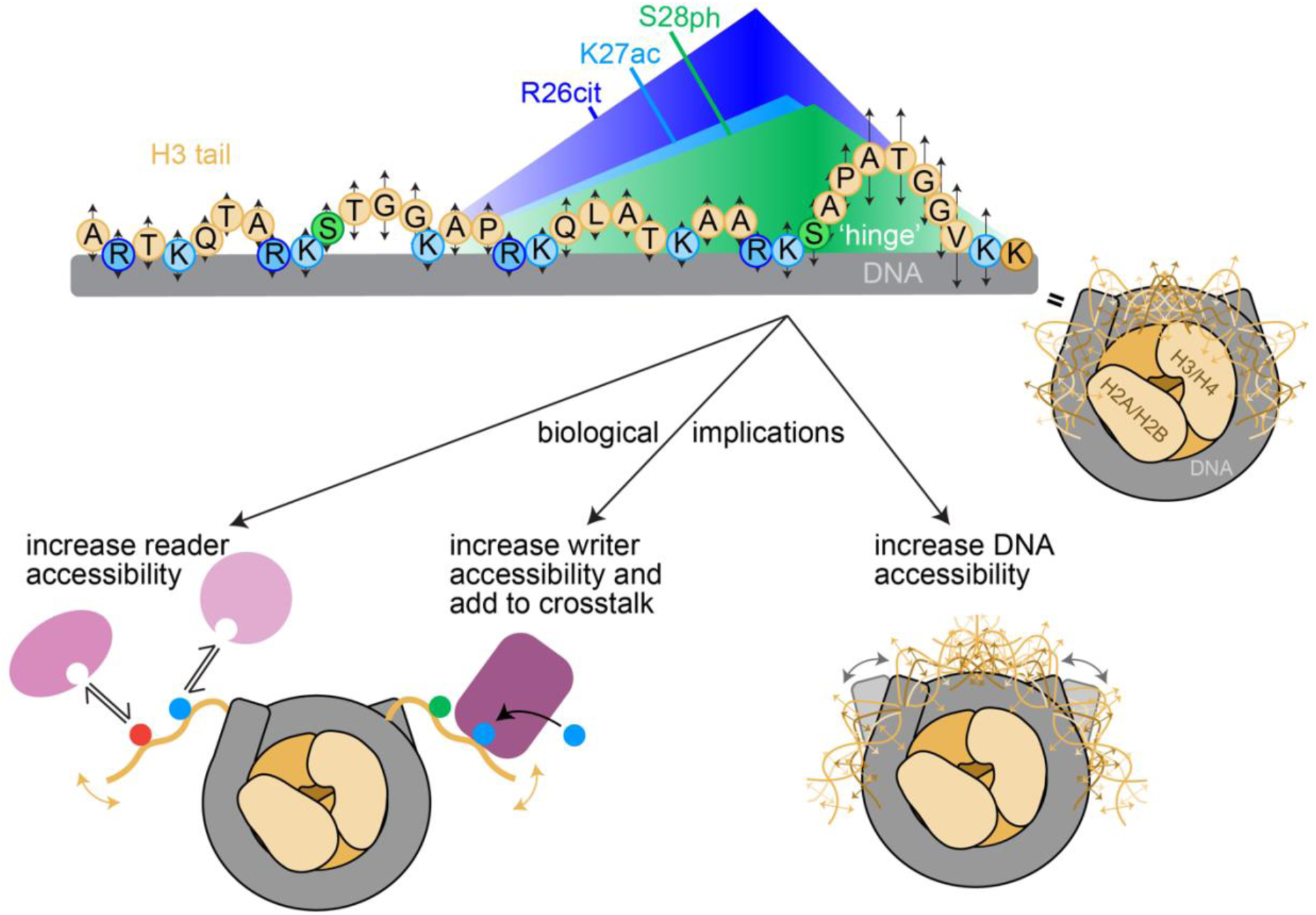
Tail dynamics in the histone language. Within the nucleosome, H3 tail residues exist in a dynamic ensemble of DNA-bound conformations (upper right). Each residue has a microscopic site-binding constant for DNA that determines the ps-ns dynamics of the residue (upper left, black arrows), and lysines and arginines have stronger site-binding constants corresponding to slower motions (shorter arrows). Charge-modulating PTMs such as arginine citrullination, lysine acetylation, and serine phosphorylation increase the regional mobility around the site of modification (depicted as a gradient triangle indicating the magnitude and breadth of effect). Implications of the increased dynamics include greater reader accessibility to the charge-modulating PTM or another nearby PTM, greater writer accessibility to nearby sites (with implications for additional crosstalk), and increased DNA dynamics and accessibility.

## MATERIALS AND METHODS

### Histone Expression and Purification

Histones were overexpressed in T7 LysY or Rosetta2 pLysS (H2B only) *E. coli* (New England Biolabs, Ipswitch, MA) that were transformed with a pET3a vector containing the histone sequence of interest (hH2A – UniProt P0C0S8; hH2B – UniProt P62807; hH3 – UniProt Q71DI3 with C110A; hH4 – UniProt P62805). H3 KQ and SE mutant histones were generated via Q5 site-directed mutagenesis (NEB), confirmed by Sanger sequencing. The ten mutant histones tested in this study were H3 K4Q, K9Q, K14Q, K18Q, K23Q, K27Q, K4/9/14/18/23/27Q (referred to as 6xKQ), S10E, S28E, and S10/28E (referred to as 2xSE). NMR-inactive histones (H2A, H2B, and H4) were grown in LB media, while ^15^N-isotopically labeled histones (WT-and mutant-H3) were grown in M9 minimal media containing ^15^N-labeled NH_4_Cl (Cambridge Isotope Laboratories, Tewksbury, MA). Expression was induced with 0.2 mM (H4) or 0.4 mM (H2A, H2B, and H3) IPTG after cultures reached an OD_600_ of 0.4. Cultures were incubated at 37°C for three (WT-H3 and H4) or four (H2A, H2B, and H3 mutants) hours post-induction, until harvest. Cell pellets were stored at -80°C until further use.

Histones were purified as previously described (Dyer et al., 2003; Paintsil & Morrison, 2022). Briefly, bacteria were lysed via sonication. Histones were extracted from inclusion bodies using a series of lysis buffers and dialyzed overnight in 8 M urea containing AmberLite MB mixed ion exchange resin (Sigma-Aldrich, St. Louis, MO) to prevent histone carbamylation. Following extraction, histones were purified over a Q Sepharose Fast Flow (Cytiva, Marlborough, MA) anion-exchange gravity chromatography column to remove protein impurities. Eluate, confirmed by 15% SDS-PAGE gels to contain histones of interest, was further purified over a Proteindex IEX-CM Agarose 6 Fast Flow (H4 only; Marvelgent Biosciences, Waltham, MA) or SP Sepharose Fast Flow (Cytiva) cation-exchange gravity chromatography column. Histones were eluted from these columns using increasing concentrations of NaCl (200-700 mM). Purified histones were progressively dialyzed into ddH_2_O containing 2 mM 2-mercaptoethanol (to maintain reduced cysteines; excluded from final dialysis), lyophilized, and stored at -20°C until further use. Electrospray ionization mass spectrometry and 15% SDS-PAGE gels were used to confirm the purity and validity of the desired histones.

### Widom 601 DNA Amplification and Purification

147-bp Widom 601 dsDNA, a strongly positioning synthetic sequence, was amplified and purified as previously described (Dyer et al., 2003; Paintsil & Morrison, 2022). This sequence is ATCGAGAATC CCGGTGCCGA GGCCGCTCAA TTGGTCGTAG ACAGCTCTAG CACCGCTTAA ACGCACGTAC GCGCTGTCCC CCGCGTTTTA ACCGCCAAGG GGATTACTCC CTAGTCTCCA GGCACGTGTC AGATATATAC ATCCGAT.

### NCP Reconstitutions

NCPs were reconstituted as previously described (Dyer et al., 2003; Hammonds & Morrison, 2022). Briefly, lyophilized histones were resuspended in a 6 M guanidine hydrochloride unfolding buffer, combined separately in equimolar ratios of H2A:H2B and ^15^N-H3(WT or mutant):H4, and dialyzed into 2 M KCl (VWR Life Sciences) to refold dimer and tetramer, respectively. Refolded dimer and tetramer were FPLC-purified over a HiLoad 16/600 Superdex 200 pg (Cytiva) size-exclusion chromatography column and then combined in a 2:1:1 or 2.2:1:1 molar ratio of dimer:tetramer:Widom 601 dsDNA. NCPs were desalted via an exponential gradient over the course of 44 hours at 4°C. Reconstituted NCPs were purified by a 10%-40% w/v sucrose gradient for 42.5 - 43 hours using an Optima XE-90 Ultracentrifuge (Beckman Coulter, Brea, CA) at 32,000 rpm and 4°C. Final nucleosomes were exchanged into NMR buffer [20 mM MOPS (with 8 mM NaOH to pH), 1 mM EDTA, 1 mM DTT, pH 7] and diluted to 100 µM NCP, including by addition of D_2_O (Cambridge Isotope Laboratories) to 5% v/v. NCP concentration was determined, following dilution into 2 M KCl, by absorbance at 260 nm using a NanoDrop (ThermoFisher, Waltham, MA) and a calculated extinction coefficient of 2,312,300 M^-1^cm^-1^ for the 147-bp Widom 601 DNA. 5% native PAGE and 18% SDS-PAGE gels were used to assess the quality of all NCPs, both before and after NMR experiments. A total of 11 NCPs were prepared for this study, one WT NCP (from (Jennings et al., 2023)) and ten containing one of the H3 mutant histones listed above.

### NMR Data Collection Parameters

All experiments were collected at 31°C (304 K) using an 18.8 T (800 MHz) Bruker (Billerica, MA) Avance Neo NMR spectrometer equipped with a cryoprobe, as previously described (Jennings et al., 2023). Four NMR experiments were conducted for each of the 11 NCPs studied: ^1^H-^15^N HSQC spectra, a ^15^N longitudinal relaxation (R_1_) series, a ^15^N transverse relaxation (R_2_) series, and a {^1^H}-^15^N steady-state heteronuclear nuclear Overhauser effect (hnNOE) series. Data for ^15^N-WT-H3-NCP are from (Jennings et al., 2023). ^1^H-^15^N HSQC spectra were used to calculate chemical shift perturbations (CSPs) between WT- and mutant-H3-NCP peaks at each residue in the H3 tail. Each ^1^H (direct dimension) FID was sampled 2048 times at an acquisition time of 98 ms. The ^15^N (indirect dimension) evolution time was incremented 400 times with an acquisition time of 82 ms. A total of 24 scans were performed for ^1^H-^15^N HSQC experiments. The three remaining NMR NSR experiments provide information on the ps-ns dynamics of the H3 tail. ^15^N-R_1_ relaxation data were collected with total relaxation loop lengths of 0.033 s, 0.195 s, 0.390 s (×2), 0.813 s (×2), 1.301 s, and 1.951 s, in randomized order. 32 scans were collected with an interscan delay of 2 s. ^15^N-R_2_ relaxation data were collected with a 500 Hz effective CPMG field and total relaxation CPMG loop lengths of 0.0088 s, 0.031 s (×2), 0.053 s, 0.079 s, 0.110 s (×2), and 0.150 s, in randomized order. A total of 32 scans were collected with an interscan delay of 2.5 s. All R_2_ experiments included temperature compensation blocks. The hnNOE pulse sequence (Bruker hsqcnoef3gps) yielded reference and saturated spectra, which were collected in an interleaved manner. 96 scans were collected with an interscan delay of 5 s. ^1^H-^15^N HSQC spectra were collected immediately prior and subsequent to all NSR experiments to confirm maintained NCP integrity throughout multi-day experiments.

### NMR Data Analysis

NMR data were processed and analyzed using NMRPipe (Delaglio et al., 1995) and CCPNMR v2.5.2 (Vranken et al., 2005) software, respectively, through NMRBox servers (Maciejewski et al., 2017) accessed with RealVNC Viewer. Spectra were processed using a cosine-squared window function for ^1^H and ^15^N dimensions. Both dimensions were doubled twice by zero-filling. Resonances were propagated into all mutant-H3 spectra from an NCP containing WT-H3 (BMRB entry 50806). Visual inspection was used to determine peak assignments. As a point of verification, our assignment of the ^15^N-S28E-H3-NCP spectra was consistent with that in Pelaz et al. (Pelaz et al., 2020). Peak assignments for NSR experiments were transferred directly from the spectrum of the highest intensity (i.e., relaxation series spectrum with the shortest delay time or hnNOE reference spectrum) to all other spectra in the respective series without further chemical shift adjustments. For each NCP, a list of doublet peaks, overlapping peaks, and questionable peak assignments is provided in **Table S1**. Due to poor signal-to-noise ratio, R2 and L20 were excluded from analysis for all NCPs.

CSPs (Δ*δ*) were calculated as √(Δ*δ_H_*)^2^ + (0.154 · Δ*δ_N_*)^2^, where Δ*δ_H_* and Δ*δ_N_* are the chemical shift differences in the ^1^H and ^15^N dimensions, respectively, for a given residue in the WT versus mutant NCP spectra, with a 0.154 weighting factor applied to the ^15^N dimension, as previously detailed (Jennings et al., 2023; Mulder et al., 1999). For relaxation series, peak intensities were fit to a single-exponential decay equation as a function of delay time, which was used to generate R_1_ and R_2_ values in CCPNMR. The R_2_/R_1_ ratio was calculated and plotted in Igor Pro (Wavemetrics, Portland, OR), with standard error propagated from the R_1_ and R_2_ covariance matrix. The reported hnNOE values for each residue were calculated as the ratio of peak intensities between the saturated and reference spectra for a given mutant-H3-NCP. Error was propagated from the baseline noise of the reference spectrum, as determined by CCPNMR, and hnNOE values were plotted as a function of H3 tail primary sequence in Igor Pro (Wavemetrics).

For each mutant-H3-NCP, the differences in R_2_/R_1_ ratios [Δ(R_2_/R_1_)] and hnNOE (ΔhnNOE) values with respect to WT-H3-NCP were determined for each residue in the H3 tail. If a given residue had doublet peaks in both the mutant and WT spectra, the Δ(R_2_/R_1_) and ΔhnNOE were calculated from peaks with minimal CSP. If a mutant NCP had a doublet for a given residue where the WT NCP had only a singlet, the Δ(R_2_/R_1_) and ΔhnNOE for each of the doublet peaks were calculated from the single WT peak. In all instances, except the 6xKQ- and 2xSE-H3-NCPs, Δ(R_2_/R_1_) and ΔhnNOE for both doublet peaks are plotted. Doublet peaks are, however, excluded from reported summations and averages to avoid biasing NCPs with more doublet peaks. Δ(R_2_/R_1_) and ΔhnNOE plots were created in Igor Pro (Wavemetrics). All figures were edited and finalized in Adobe Illustrator (San Jose, CA).

## DATA AVAILABILITY

The data underlying this article are available in the manuscript, the supplementary materials, and upon reasonable request. Plasmids are available upon reasonable request. The relaxation datasets presented in this study are deposited in the Biological Magnetic Resonance Data Bank (BMRB) with deposition numbers 53858, 53859, 53860, 53861, 53862, 53863, 53864, 53865, 53866, and 53867.

## SUPPLEMENTARY MATERIALS

Supplementary figures and tables that show NMR spectra, additional data plots, and tabulated nuclear spin relaxation data are available online in a document entitled “SupplementaryMaterial_AdkinsEtAl.pdf”.

## ACKNOWLEDGMENTS

Thanks to Drs. Catherine Musselman, Karolin Luger, and Michael Poirier for gifts of histone plasmids. Thank you to Dr. Ananya Majumdar for the ^15^N-R_1_ and ^15^N-R_2_ pulse sequences. Thank you to Alex Kowalczyk for helpful edits to this manuscript. Thank you to Sarah Meidl Zahorodny for purification of histones and DNA.

## AUTHOR CONTRIBUTIONS

E.M. conceived of the initial project. B.A. and E.M. designed the studies. E.M. acquired funding and supervised the research. B.A., P.S., and C.J. prepared samples and performed the research. B.A. analyzed and visualized the data. B.A. and E.M. wrote the manuscript.

## MATERIALS & CORRESPONDENCE

Correspondence and material requests should be addressed to Emma Morrison at.

