## Supplementary Materials for "Modulating Nucleosomal H3 Tail Dynamics with Lysine and Serine Modifications"

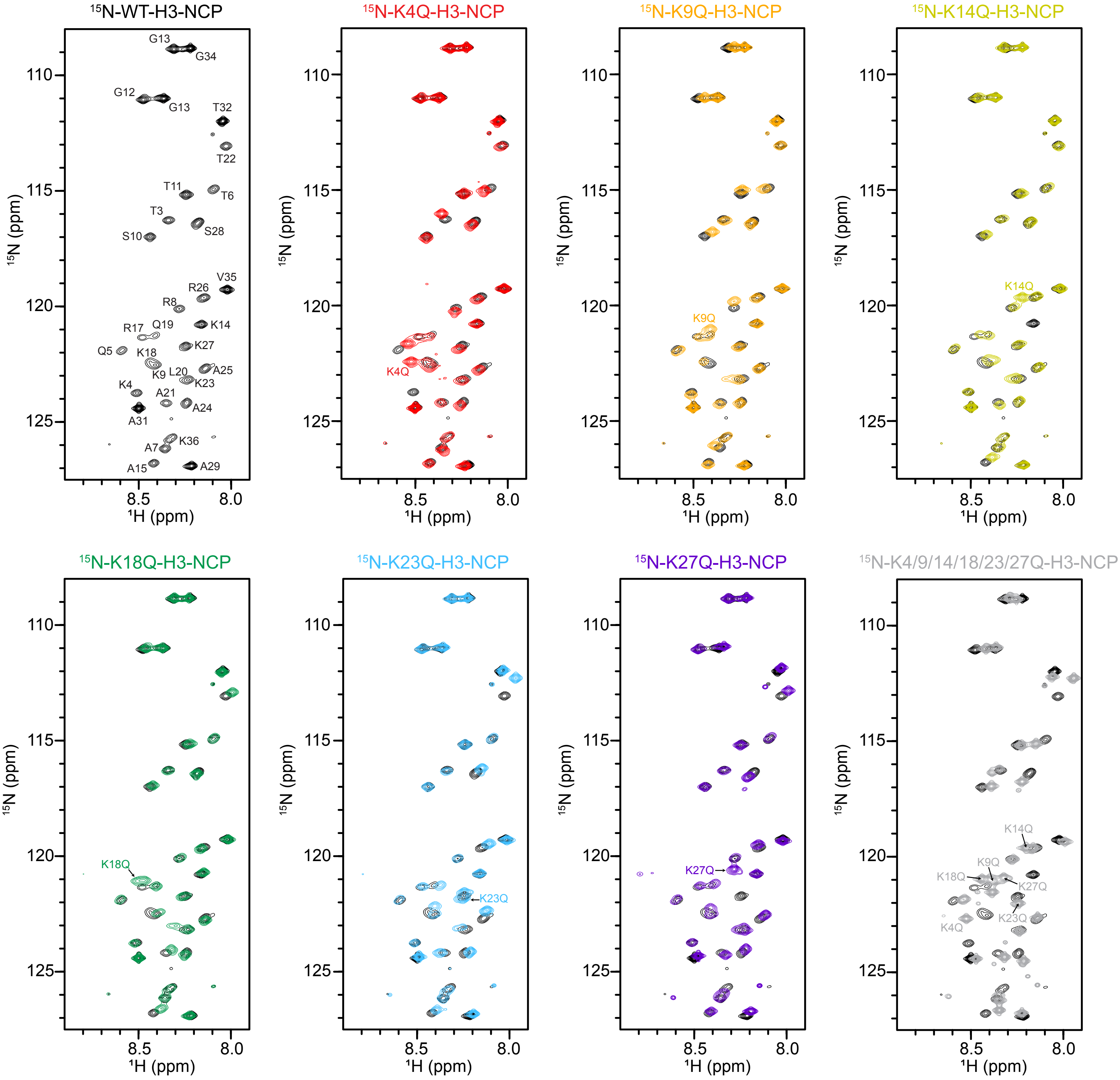


**Figure S1.** Overlays of 1H-15N HSQC spectra of 15N-WT-H3-NCP with each 15N-KQ-H3-NCP. Spectra are colored to match the main text figure color scheme and displayed at the same contour. Assignments are labeled in the WT-H3-NCP spectrum, and lysine-to-glutamine mutations are labeled in overlays. Data were collected at 800 MHz and 304 K.

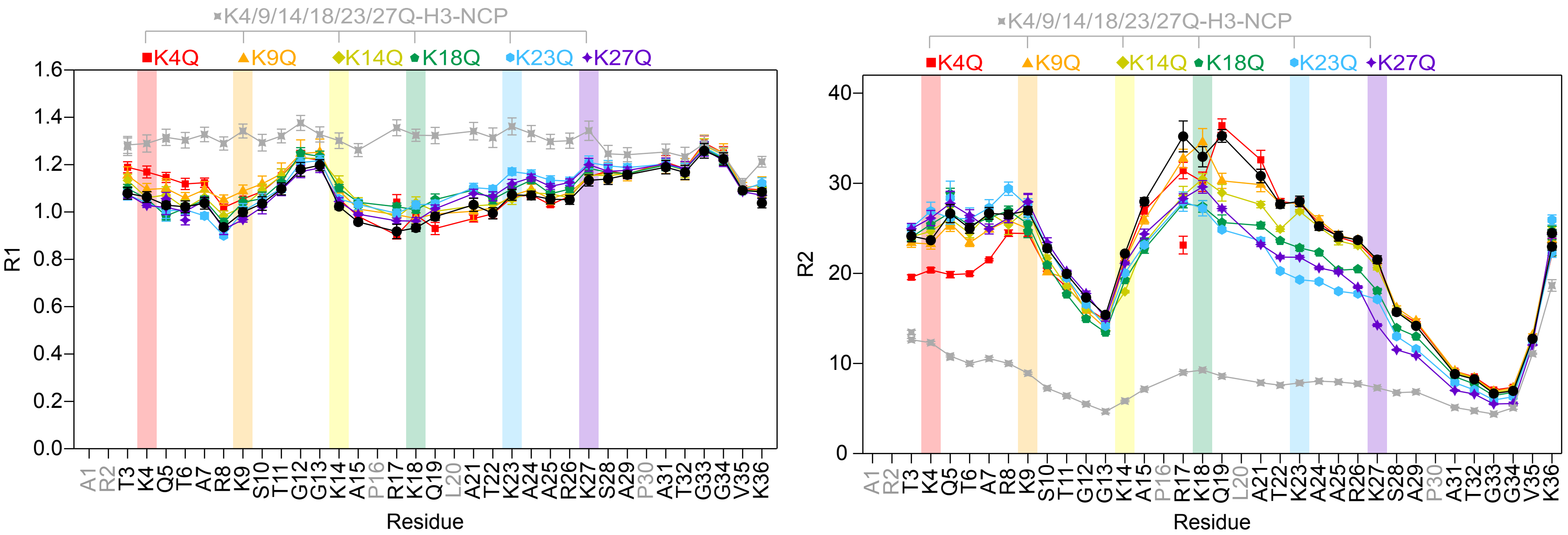


**Figure S2.** Plot of 15N-R1 rates (left) and 15N-R2 rates (right) as a function of residue for K4Q- (red squares), K9Q- (orange triangles), K14Q- (yellow diamonds), K18Q- (green pentagons), K23Q- (blue hexagons), K27Q- (purple stars), and K4/9/14/18/23/27Q-H3-NCP (gray stars). Error bars were determined via the covariance matrix in fitting R1 and R2 decay curves. Note that some error bars are smaller than the symbols. Residues without data for any sample are colored grey in the x-axis labels. Data were collected at 800 MHz and 304 K.

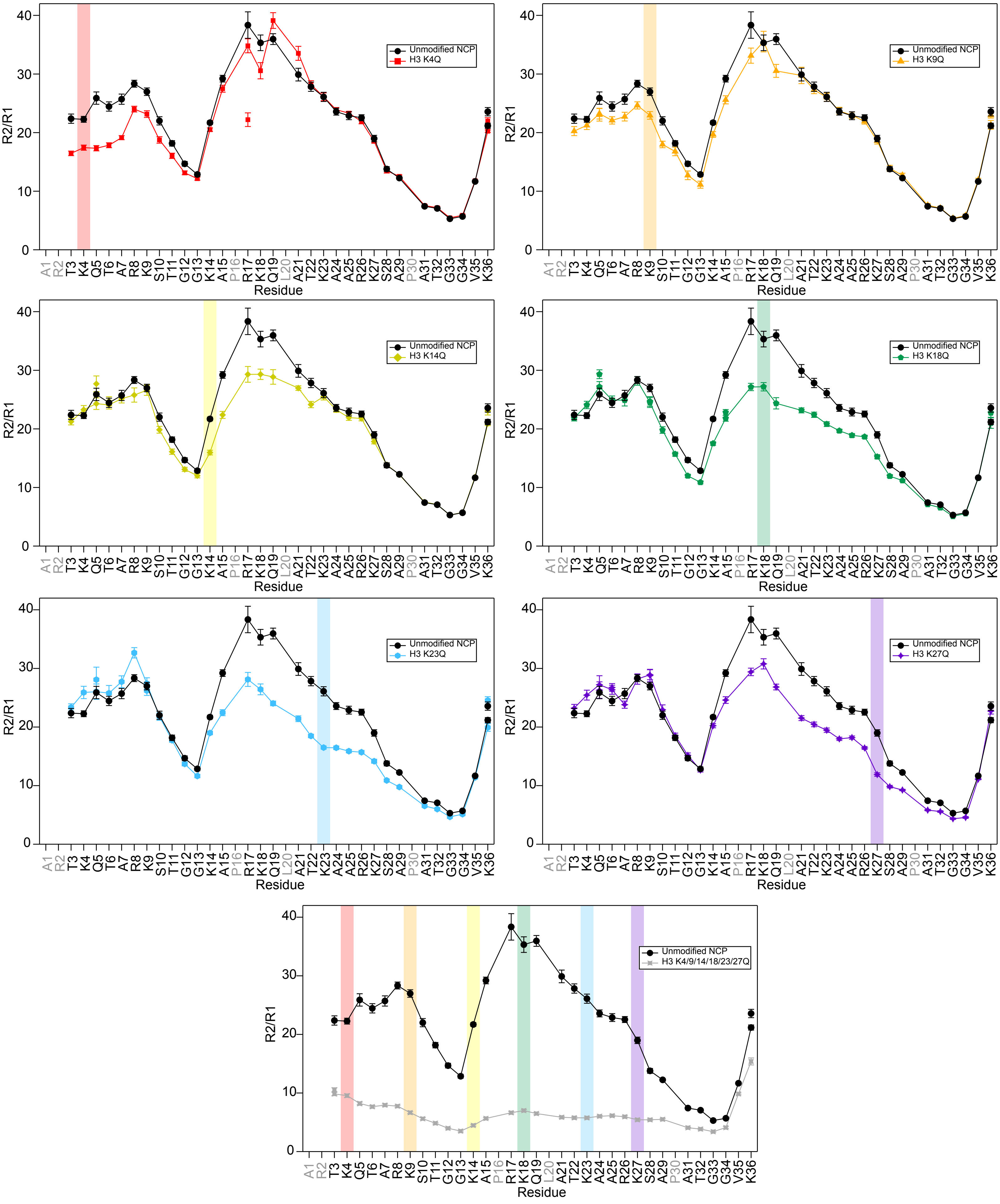


**Figure S3.** Amide nuclear spin relaxation data is shown for each 15N-KQ-H3-NCP plotted with WT (black circles) for a direct comparison. Plots of R2/R1 as a function of H3 tail residue for K4Q- (red squares), K9Q- (orange triangles), K14Q- (yellow diamonds), K18Q- (green pentagons), K23Q- (blue hexagons), K27Q- (purple stars), and K4/9/14/18/23/27Q-H3-NCP (gray stars). Mutated positions are marked by a colored background bar. Error bars represent standard error propagation from the covariance matrix in fitting R1 and R2 decay curves. Note that some error bars are smaller than the symbols. Residues without data for any sample are colored grey in the x-axis labels. Data were collected at 800 MHz and 304 K.

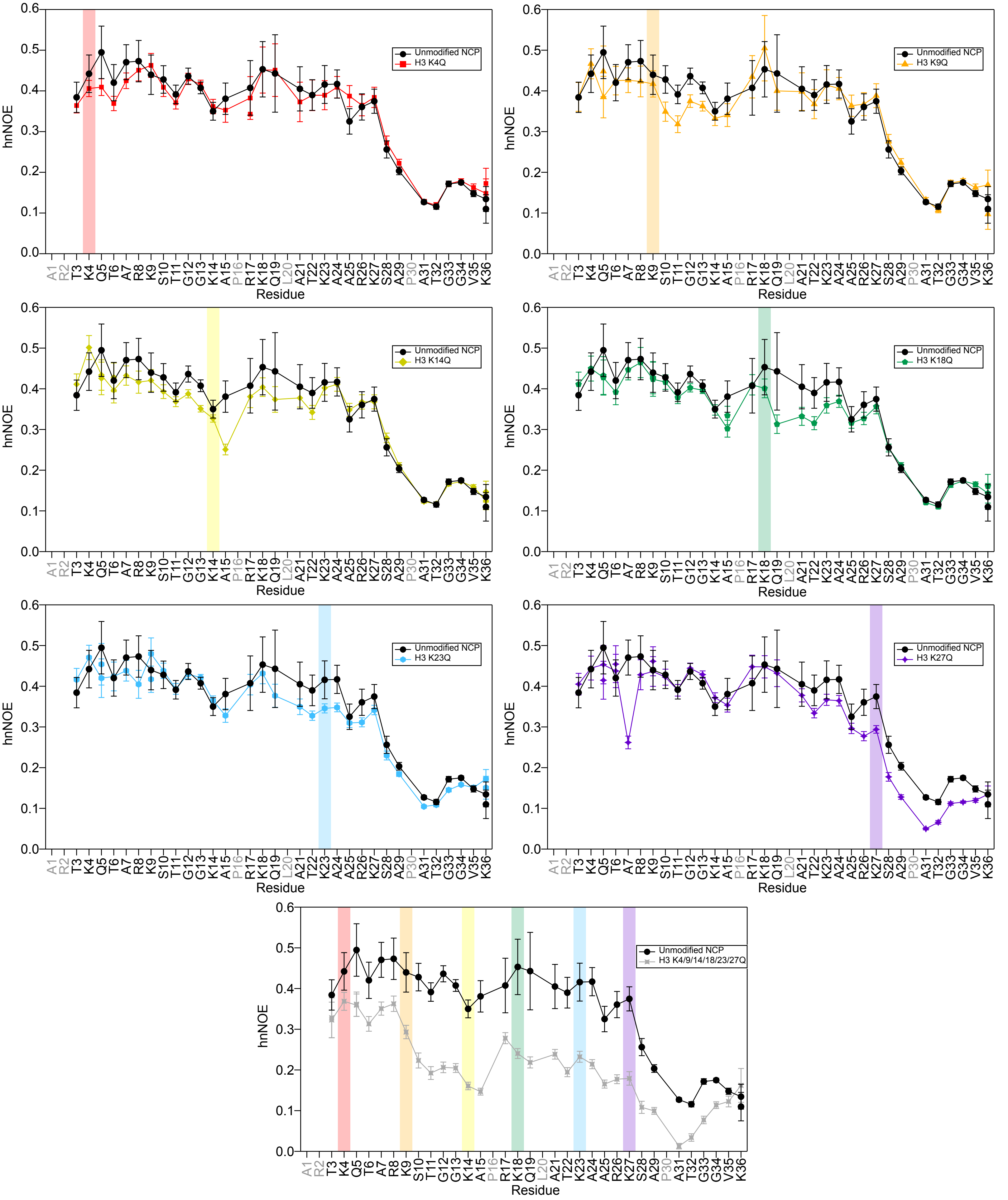


**Figure S4.** Amide nuclear spin relaxation data is shown for each 15N-KQ-H3-NCP plotted with WT (black circles) for a direct comparison. Plots of hnNOE values as a function of H3 tail residue for K4Q- (red squares), K9Q- (orange triangles), K14Q- (yellow diamonds), K18Q- (green pentagons), K23Q- (blue hexagons), K27Q- (purple stars), and K4/9/14/18/23/27Q-H3-NCP (gray stars). Mutated positions are marked by a colored background bar. Error bars represent standard error propagation of the spectral noise for hnNOE values. Note that some error bars are smaller than the symbols. Residues without data for any sample are colored grey in the x-axis labels. Data were collected at 800 MHz and 304 K.

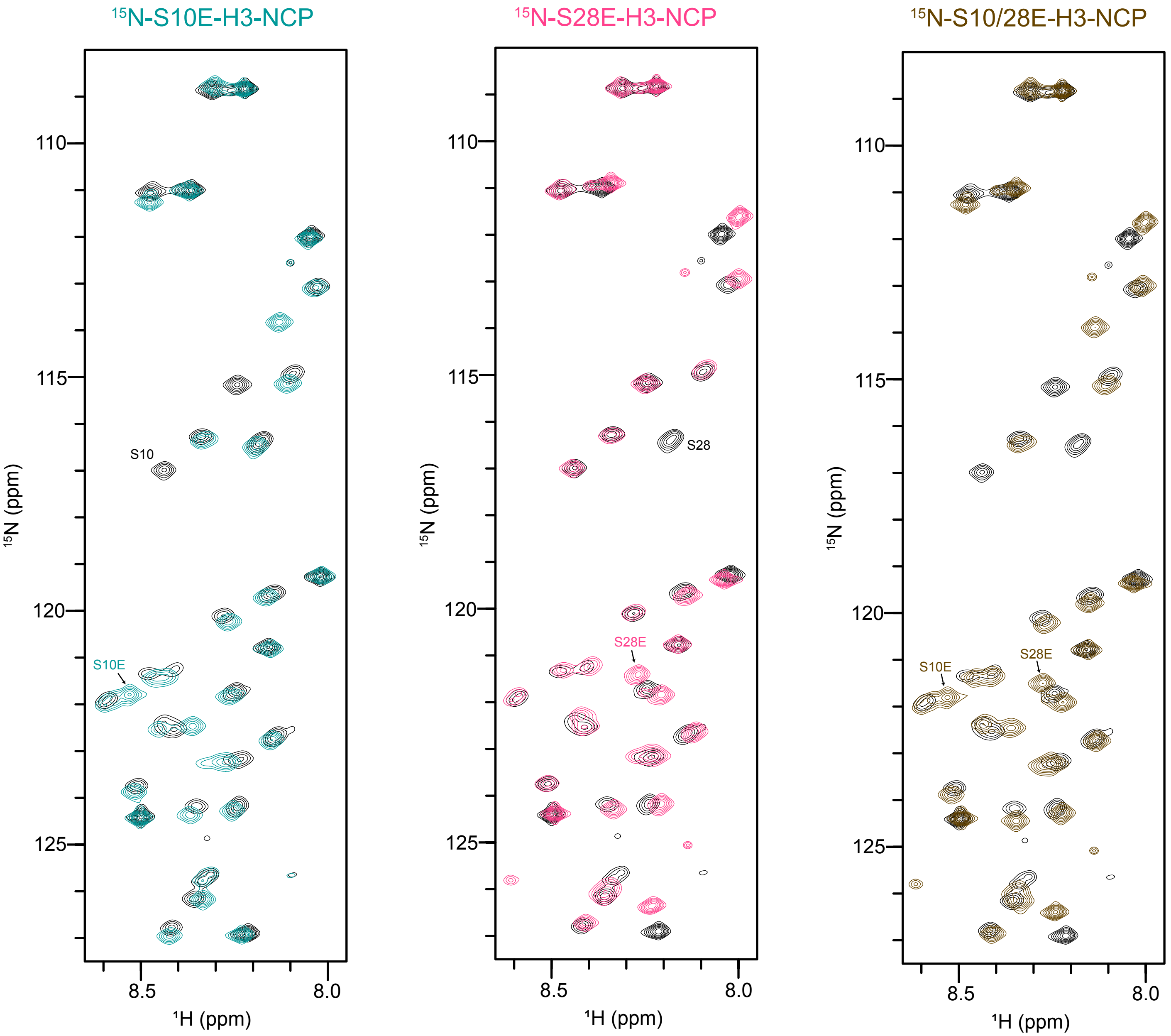


**Figure S5.** Overlays of 1H-15N HSQC spectra of 15N-WT-H3-NCP with each 15N-SE-H3-NCP. Spectra are colored to match the main text figure color scheme and displayed at the same contour. Assignments of serine-to-glutamate mutations are labeled in overlays. Data were collected at 800 MHz and 304 K. A complete labeling of 15N-WT-H3-NCP assignments can be found in Figure S2.

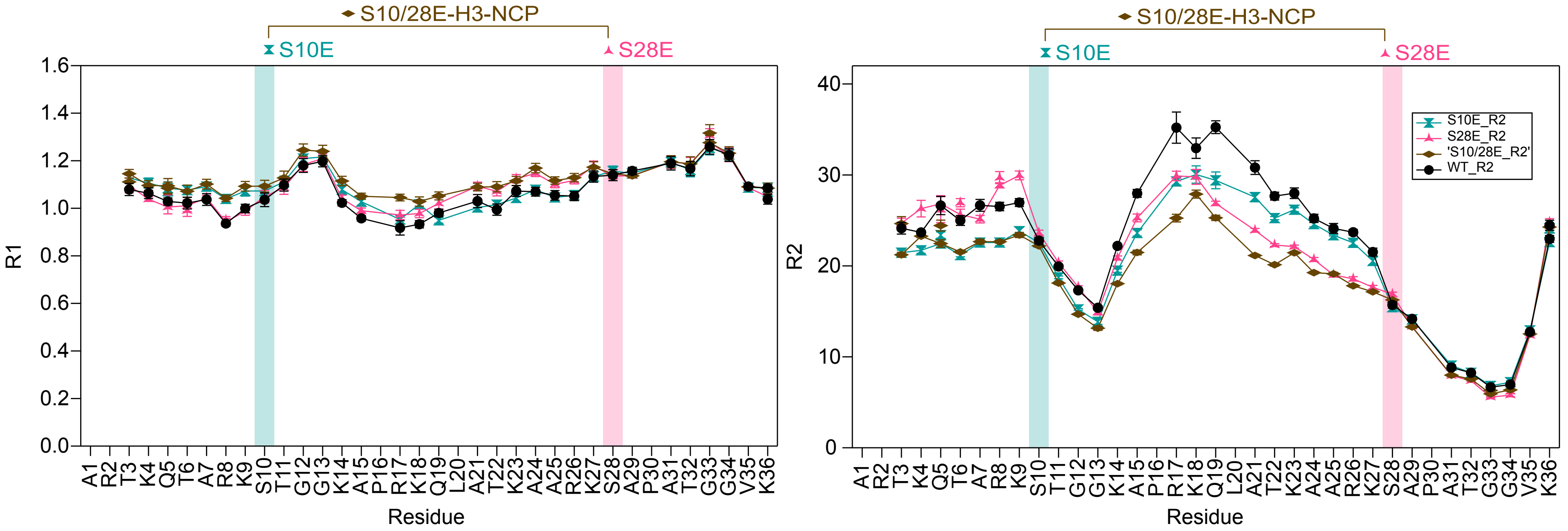


**Figure S6.** Plot of 15N-R1 rates (left) and 15N-R2 rates (right) as a function of H3 tail residue for S10E- (cyan hourglasses), S28E- (magenta arrowheads), and S10/28E-H3-NCP (brown diamonds). Error bars were determined via the covariance matrix in fitting R1 and R2 decay curves. Note that some error bars are smaller than the symbols. Residues without data for any sample are colored grey in the x-axis labels. Data were collected at 800 MHz and 304 K.

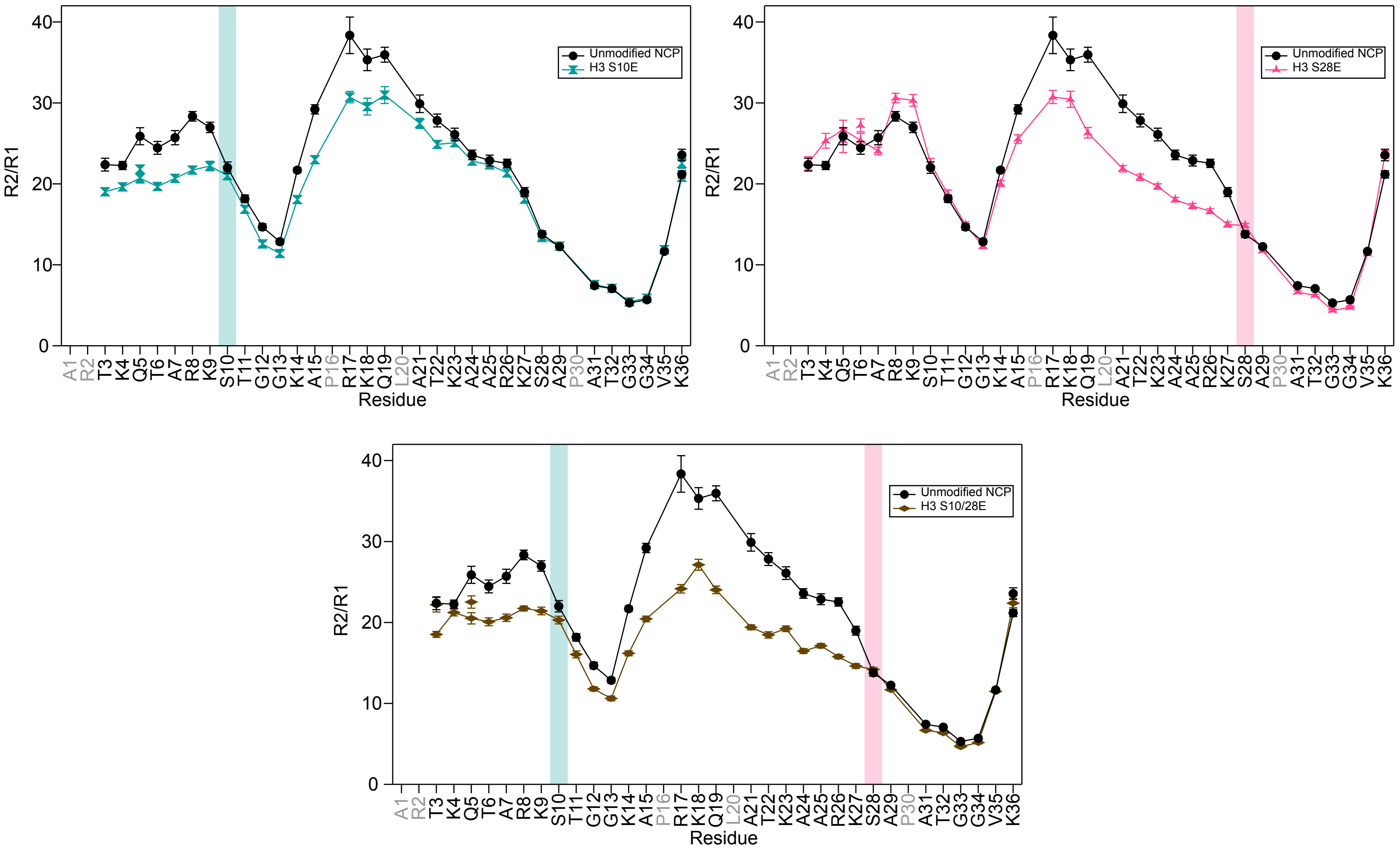


**Figure S7.** Amide nuclear spin relaxation data is shown for each 15N-SE-H3-NCP plotted with WT (black circles) for a direct comparison. Plots of R2/R1 as a function of H3 tail residue for S10E- (cyan hourglasses), S28E- (magenta arrowheads), and S10/28E-H3-NCP (brown diamonds). Mutated positions are marked by a colored background bar. Error bars represent standard error propagation from the covariance matrix in fitting R1 and R2 decay curves. Note that some error bars are smaller than the symbols. Residues without data for any sample are colored grey in the x-axis labels. Data were collected at 800 MHz and 304 K.

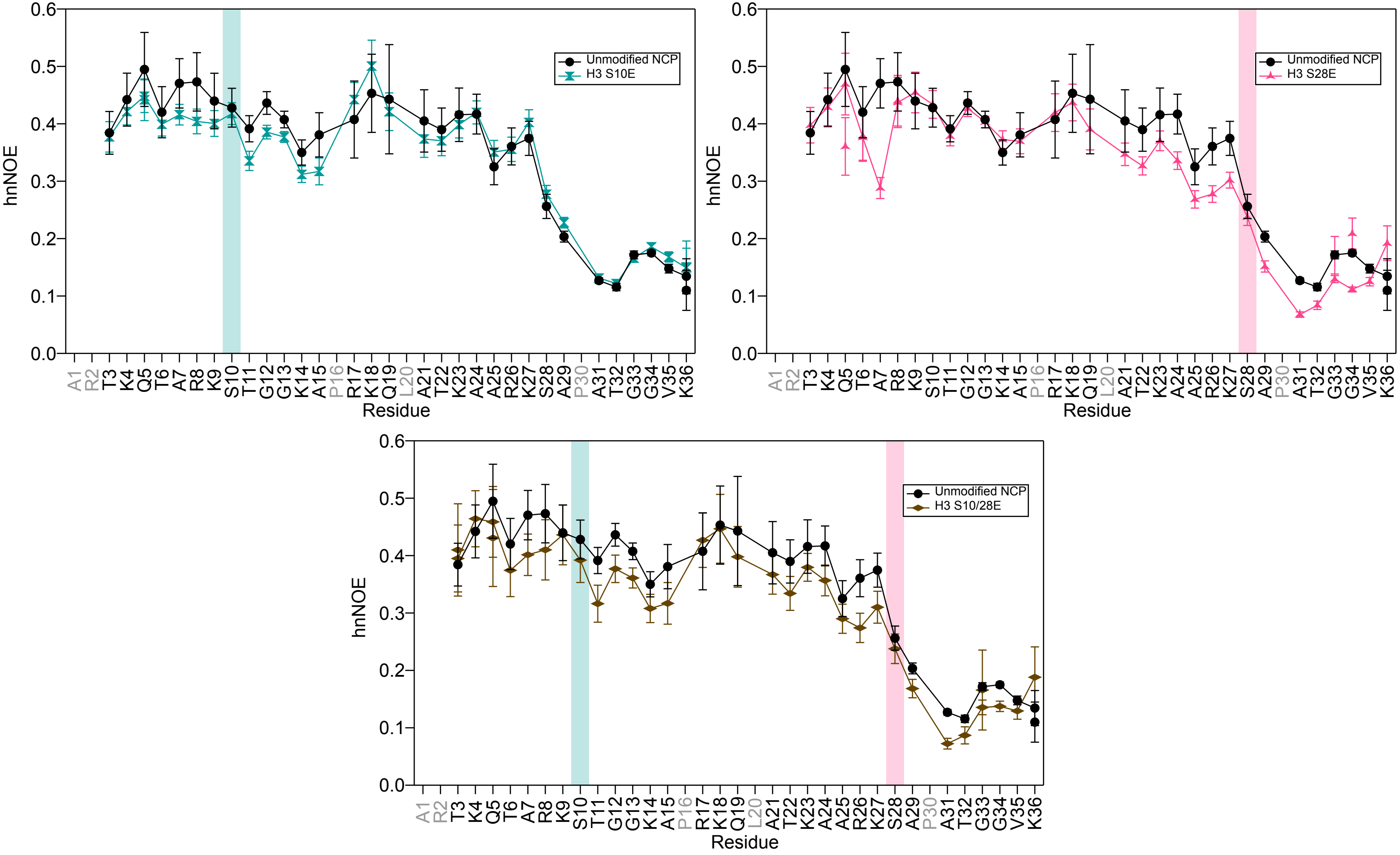


**Figure S8.** Amide nuclear spin relaxation data is shown for each 15N-SE-H3-NCP plotted with WT (black circles) for a direct comparison. Plots of hnNOE values as a function of H3 tail residue for S10E- (cyan hourglasses), S28E- (magenta arrowheads), and S10/28E-H3-NCP (brown diamonds). Mutated positions are marked by a colored background bar. Error bars represent standard error propagation of the spectral noise for hnNOE values. Note that some error bars are smaller than the symbols. Residues without data for any sample are colored grey in the x-axis labels. Data were collected at 800 MHz and 304 K.

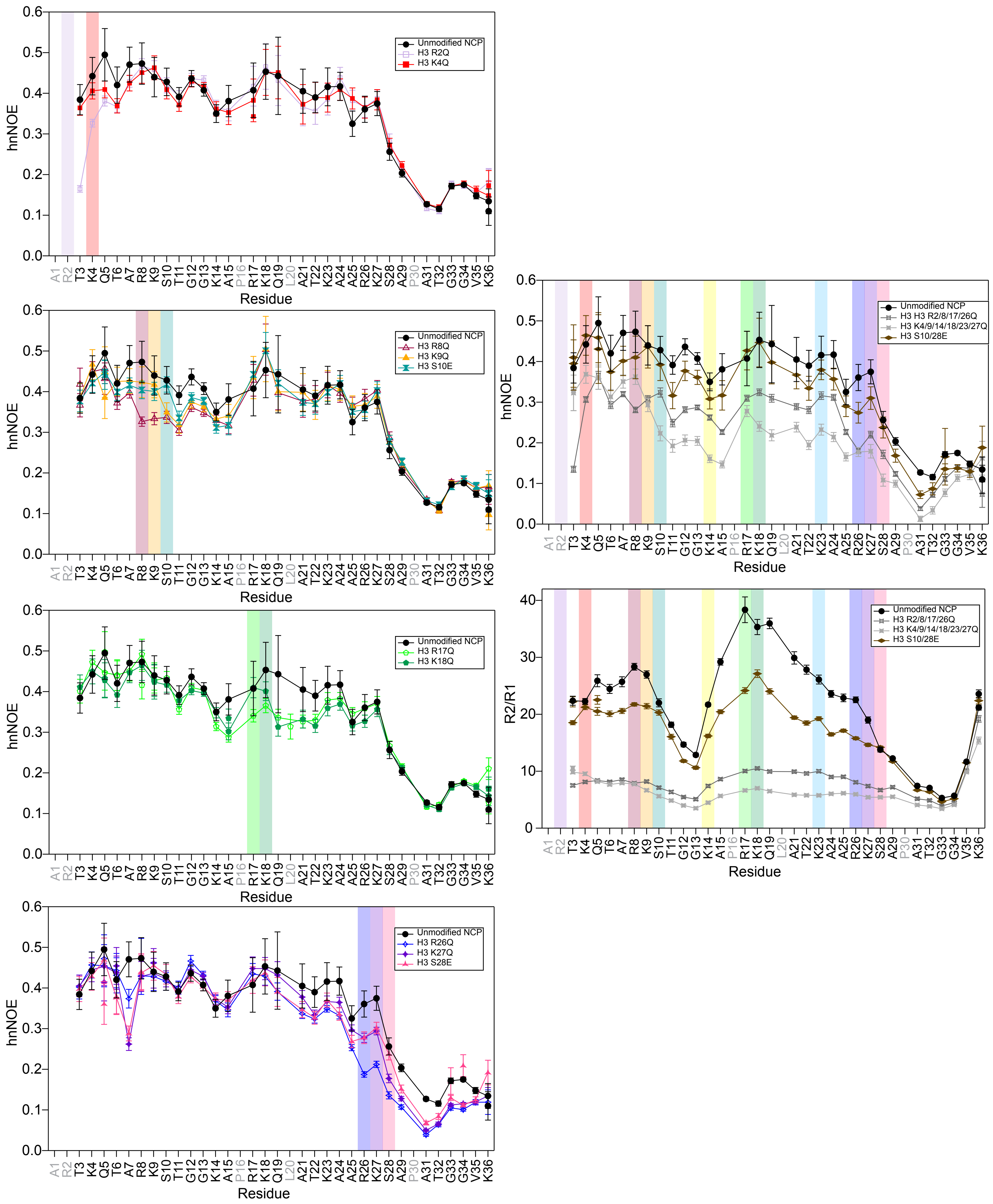


**Figure S9.** Amide nuclear spin relaxation data collected on 15N-H3-NCP is directly compared for nearby charge-modulating mutations of lysine, serine, and arginine. Mutated positions are marked by a colored background bar. Error bars represent standard error propagation of the spectral noise for hnNOE values. Note that some error bars are smaller than the symbols. Residues without data for any sample are colored grey in the x-axis labels. Data were collected at 800 MHz and 304 K. Data from arginine-to-glutamine mutations are from Jennings, et al.

**Supplementary Table S1.** Summary of NCP sample spectral commentary on residues showing doublets and overlap, and with lower certainty assignment transfer. Data from arginine-to-glutamine mutations are from Jennings et al., 2023.

| NCP | Doublets | Peaks with Overlap | Lower Certainty Assignments |
|---|---|---|---|
| WT | K36 | K9, R17, K18, Q19, L20, K23 | None |
| K4Q-H3 | R17, K36 | K9, R17, K18, Q19, L20, K23 | None |
| K9Q-H3 | Q5, K36 | A7, K9Q, G13, R17, Q19, L20, K23, G34, K36 | K9Q, R17, Q19 |
| K14Q-H3 | Q5, T6, K36 | A7, R8, K9, K14Q, R17, K18, Q19, L20, K23, R26, K36 | K9, K18 |
| K18Q-H3 | Q5, K9, A15, K36 | A7, A15, R17, K18Q, L20, K23, K36 | R17, K18Q |
| K23Q-H3 | Q5, K9, K36 | A7, K9, K18, K23Q, K27, K36 | K23Q, K27 |
| K27Q-H3 | Q5, T6, K9 | A7, R8, K9, R17, K18, Q19, L20, K23, K27Q, K36 | None |
| 6xKQ-H3 | T3, Q5 | A7, K9Q, G12, G13, K14Q, R17, K18Q, Q19, L20, K23Q, R26, K27Q, G33, G34, K36 | K9Q, R17, K18Q, Q19, K27Q |
| S10E-H3 | Q5, K36 | Q5, S10E, R17, Q19, L20, K23 | None |
| S28E-H3 | Q5, T6, R8, G33, G34 | A7, K9, G13, K18, L20, K23, K36 | None |
| 2xSE-H3 | T3, Q5, G33, G34, K36 | Q5, A7, K9, S10E, G13, R17, K18, Q19, L20, K23, G34, K36 | None |
| R2Q-H3 | K36 | K9, R17, K18, Q19, L20, K23, K36 | None |
| R8Q-H3 | R2, T3, Q5, K36 | T3, Q5, A7, R17, Q19, L20, K23, K36 | None |
| R17Q-H3 | R2, Q5, T6, R8, K9, L20, K36 | Q5, T6, A7, R8, K9, K18, L20, K23, K36 | K9, K18, L20, K23 |
| R26Q-H3 | R2, Q5, T6, R8 | Q5, T6, A7, R8, K9, K18, L20, K23, K36 | None |
| 4xRQ-H3 | No doublets | K9, K18, K36 | L20, K23 |

**Supplementary Table S2.** Compilation of average (including standard deviation), minimum, and maximum values for hnNOE, R2/R1, R1, and R2 for each WT or mutant 15N-H3-NCP at 0 mM KCl. Values are listed for all analyzed residues in the H3 tail, without doublets included in the analysis (as not to bias toward NCPs with the most doublets). See Materials and Methods for additional details. Data from WT and arginine-to-glutamine mutations are from Jennings et al., 2023, but were reanalyzed to uniformly exclude doublets and L20 for a consistent comparison.

| $R_2/R_1$ Ratio | | | | | | | | | | | | | | | | |
|---|---|---|---|---|---|---|---|---|---|---|---|---|---|---|---|---|
| | WT | K4Q | K9Q | K14Q | K18Q | K23Q | K27Q | 6xKQ | S10E | S28E | 2xSE | R2Q | R8Q | R17Q | R26Q | 4xRQ |
| Minimum | 5 | 5 | 5 | 5 | 5 | 5 | 4 | 3 | 5 | 4 | 5 | 5 | 5 | 5 | 4 | 4 |
| Average | 21 ± 9 | 20 ± 8 | 20 ± 8 | 19 ± 7 | 18 ± 7 | 18 ± 8 | 19 ± 8 | 6 ± 2 | 19 ± 7 | 19 ± 8 | 17 ± 6 | 19 ± 9 | 17 ± 8 | 17 ± 6 | 17 ± 8 | 8 ± 3 |
| Maximum | 38 | 39 | 36 | 29 | 28 | 33 | 31 | 15 | 31 | 31 | 27 | 37 | 33 | 27 | 31 | 19 |

| $R_1$ ($s^{-1}$) | | | | | | | | | | | | | | | | |
|---|---|---|---|---|---|---|---|---|---|---|---|---|---|---|---|---|
| | WT | K4Q | K9Q | K14Q | K18Q | K23Q | K27Q | 6xKQ | S10E | S28E | 2xSE | R2Q | R8Q | R17Q | R26Q | 4xRQ |
| Minimum | 0.92 | 0.90 | 0.97 | 0.98 | 0.96 | 0.90 | 0.93 | 1.12 | 0.95 | 0.94 | 1.03 | 0.93 | 0.95 | 0.99 | 0.90 | 1.11 |
| Average | 1.07 ± 0.09 | 1.1 ± 0.1 | 1.11 ± 0.08 | 1.1 ± 0.08 | 1.1 ± 0.08 | 1.1 ± 0.09 | 1.09 ± 0.09 | 1.3 ± 0.05 | 1.09 ± 0.08 | 1.09 ± 0.08 | 1.12 ± 0.06 | 1.1 ± 0.1 | 1.12 ± 0.09 | 1.13 ± 0.08 | 1.12 ± 0.1 | 1.26 ± 0.05 |
| Maximum | 1.26 | 1.29 | 1.29 | 1.28 | 1.27 | 1.27 | 1.27 | 1.38 | 1.25 | 1.28 | 1.28 | 1.27 | 1.27 | 1.28 | 1.29 | 1.36 |

| $R_2$ ($s^{-1}$) | | | | | | | | | | | | | | | | |
|---|---|---|---|---|---|---|---|---|---|---|---|---|---|---|---|---|
| | WT | K4Q | K9Q | K14Q | K18Q | K23Q | K27Q | 6xKQ | S10E | S28E | 2xSE | R2Q | R8Q | R17Q | R26Q | 4xRQ |
| Minimum | 7 | 7 | 7 | 7 | 6 | 6 | 5 | 4 | 7 | 6 | 6 | 7 | 7 | 6 | 5 | 5 |
| Average | 22 ± 8 | 21 ± 7 | 21 ± 7 | 21 ± 7 | 20 ± 7 | 20 ± 7 | 20 ± 7 | 8 ± 3 | 20 ± 7 | 20 ± 7 | 18 ± 6 | 20 ± 8 | 19 ± 7 | 19 ± 6 | 18 ± 8 | 10 ± 3 |
| Maximum | 35 | 36 | 35 | 30 | 28 | 29 | 30 | 19 | 30 | 30 | 28 | 34 | 31 | 27 | 28 | 22 |

| hnNOE | | | | | | | | | | | | | | | | |
|---|---|---|---|---|---|---|---|---|---|---|---|---|---|---|---|---|
| | WT | K4Q | K9Q | K14Q | K18Q | K23Q | K27Q | 6xKQ | S10E | S28E | 2xSE | R2Q | R8Q | R17Q | R26Q | 4xRQ |
| Minimum | 0.12 | 0.12 | 0.11 | 0.12 | 0.11 | 0.10 | 0.05 | 0.01 | 0.12 | 0.07 | 0.07 | 0.11 | 0.10 | 0.12 | 0.04 | 0.04 |
| Average | 0.35 ± 0.12 | 0.34 ± 0.11 | 0.34 ± 0.11 | 0.33 ± 0.11 | 0.33 ± 0.11 | 0.33 ± 0.11 | 0.32 ± 0.13 | 0.21 ± 0.09 | 0.34 ± 0.11 | 0.32 ± 0.12 | 0.31 ± 0.11 | 0.34 ± 0.11 | 0.32 ± 0.11 | 0.33 ± 0.11 | 0.3 ± 0.14 | 0.23 ± 0.09 |
| Maximum | 0.49 | 0.46 | 0.50 | 0.50 | 0.46 | 0.48 | 0.45 | 0.37 | 0.50 | 0.47 | 0.46 | 0.47 | 0.50 | 0.49 | 0.47 | 0.37 |

**Supplementary Table S3.** Compilation of average, standard deviation, and sum of all values for Δ(R2/R1) and ΔhnNOE (either raw or absolute values) for each mutant 15N-H3-NCP as compared to WT. Values exclude doublets from the analysis (as not to bias toward the NCPs with the most doublets). See Materials and Methods for additional details. Data from WT and arginine-to-glutamine mutations are from Jennings et al., 2023, but were reanalyzed to uniformly exclude doublets and L20 for a consistent comparison.

| $\Delta R_2/R_1$ | | | | | | | | | | | | | | |
|---|---|---|---|---|---|---|---|---|---|---|---|---|---|---|
| | K4Q | K9Q | K14Q | K18Q | K23Q | K27Q | 6xKQ | S10E | S28E | 2xSE | R2Q | R8Q | R17Q | R26Q | 4xRQ |
| Average | 2 | 1 | 2 | 3 | 3 | 2 | 15 | 2 | 2 | 5 | 2 | 4 | 4 | 4 | 13 |
| Standard Deviation | 3 | 2 | 2 | 3 | 4 | 4 | 8 | 2 | 4 | 4 | 4 | 4 | 5 | 5 | 8 |
| Sum (raw values) | 53 | 44 | 57 | 90 | 94 | 77 | 462 | 76 | 61 | 142 | 68 | 126 | 130 | 136 | 409 |
| Sum (absolute values) | 71 | 48 | 60 | 97 | 120 | 102 | 462 | 78 | 92 | 145 | 82 | 127 | 140 | 154 | 409 |

| ΔhnNOE | | | | | | | | | | | | | | | |
|---|---|---|---|---|---|---|---|---|---|---|---|---|---|---|---|
| | K4Q | K9Q | K14Q | K18Q | K23Q | K27Q | 6xKQ | S10E | S28E | 2xSE | R2Q | R8Q | R17Q | R26Q | 4xRQ |
| Average | 0.01 | 0.01 | 0.02 | 0.02 | 0.02 | 0.03 | 0.14 | 0.01 | 0.03 | 0.04 | 0.01 | 0.02 | 0.02 | 0.05 | 0.12 |
| Standard Deviation | 0.03 | 0.04 | 0.04 | 0.03 | 0.03 | 0.05 | 0.06 | 0.03 | 0.04 | 0.03 | 0.05 | 0.05 | 0.04 | 0.05 | 0.05 |
| Sum (raw values) | 0.23 | 0.37 | 0.62 | 0.74 | 0.61 | 1.02 | 4.47 | 0.37 | 1.03 | 1.09 | 0.46 | 0.69 | 0.55 | 1.40 | 3.58 |
| Sum (absolute values) | 0.6 | 0.9 | 0.95 | 0.88 | 0.99 | 1.29 | 4.52 | 0.9 | 1.3 | 1.30 | 1.00 | 1.23 | 1.02 | 1.70 | 3.58 |
